# Unlocking a flexible set of phylogenetic models for discrete and continuous trait evolution using discretized stochastic diffusion

**DOI:** 10.64898/2026.04.20.719455

**Authors:** Liam J. Revell, Laura R. V. Alencar, Michael E. Alfaro, Jonathan Dain, Nichola J. Hill, Menna Jones, Kristen M. Martinet, Viviana Romero-Alarcon, Luke J. Harmon

**Affiliations:** Department of Biology, University of Massachusetts Boston; Department of Ecology and Evolutionary Biology, Yale University; Department of Ecology and Evolutionary Biology, University of California, Los Angeles; Department of Geophysical Sciences, University of Chicago; College of Information Science, University of Arizona; Department of Biological Sciences, University of Idaho

## Abstract

The practical utility of many modern phylogenetic comparative methods can depend on how accurately mathematical models capture the evolutionary process of traits. Boucher and Démery (2016) described a new quantitative trait model for testing hypotheses about constraint on phenotypic character evolution: Brownian motion with reflective limits. Since their analytic solution for the probability function under this bounded scenario was intractable for reasonably-sized trees, Boucher and Démery (2016) also identified a creative technique for computing the likelihood of their model. The basis of this method derives from the convergence of an equal-rates, symmetric, ordered Markov chain and continuous stochastic diffusion in the limit as the number of lattice states in the chain goes to *∞* (or, alternatively, as their widths decrease towards zero). We realized that this general approach had the potential to unlock a surprisingly large number of additional models for the phylogenetic comparative analysis of discrete and continuous trait data, and we explore several of these in the present article. Specifically, we examine application of this ‘discretized diffusion approximation’ to the threshold model from evolutionary quantitative genetics, to a new “semi-threshold” trait evolution model, to a joint model where the rate of evolution for a continuous trait depends on the value of a co-evolving discrete character, along with a separate model where precisely the converse is true, and to a discrete-character-dependent multi-trend trended continuous trait evolution model. We conclude with some context for the origins of our article and discussion of other possible applications of this powerful approach.

## 1 Introduction

Over a decade ago an article was published by two researchers in the journal *Systematic Biology* describing a model for trait evolution that the authors referred to as “bounded Brownian motion,” along with procedures to fit this model to data (Boucher and Démery 2016). Modeling the evolutionary process as bounded might capture trait evolution on a flat adaptive landscape with hard constraints, among other possible evolutionary phenomena (Boucher and Démery 2016). Though well-received, this paper has been relatively poorly-cited. At the time of initial submission of our article, for example, Boucher and Démery (2016) had been cited a total of 55 times over the decade or so since it was originally published, according to Google Scholar (and this includes a conspicuous surge of new citations that the paper has accumulated since one of us, LR, started enthusiastically blogging and speaking about its importance in 2024). A subsequent article by Boucher et al. (2018) substantively extends Boucher and Démery (2016) by providing a much more general and flexible framework for the analysis of continuous trait evolution; however, we believe that this article, too, is underappreciated, having been cited a mere 52 times, as of the time of writing. A common feature of both articles is a creative procedure that the authors identified to compute the probability of our data under these different, novel models in phylogenetic comparative biology, where analytic functions were unavailable or impractical to evaluate for even relatively modestly sized phylogenies (Boucher and Démery 2016; Boucher et al. 2018).

The fundamental purpose of our article is to describe the powerful and creative estimation procedure used by Boucher and Démery (2016) and Boucher et al. (2018) – an approach that ultimately derives from the mathematical and physical sciences literature (e.g., see Ethier and Kurtz 1986; Risken 1996; Kushner and Dupuis 2001; Gardiner 2009), but that was totally novel in phylogenetic comparative research. We’ll then examine multiple new applications of this estimation technique, including using it to fit wholly or partially new models for phylogenetic comparative biology without tractable or closed form solutions for the likelihood function. We’ve subdivided our article, which is essentially a concept piece, into three main sections. The first and current section (labeled **Introduction**) is intentionally pedagogic and will attempt to explain, in plain language, and numerically illustrate the estimation technique of Boucher and Démery (2016). The next section (labeled **Applications**) will overview several new phylogenetic analyses that this technique enables, focusing on five specific models already implemented in software, at least four of which are partly or wholly novel to this article. Lastly, the final section (labeled **Concluding thoughts**) will provide some additional context by discussing the origins and background for this article, as well as numerous possible future directions.

The major thesis of our article is that the estimation technique of Boucher and Démery (2016) is a flexible and powerful method that will allow us to fit comparative data to a number of interesting new phylogenetic models presently lacking closed form or tractable solutions for the likelihood function, and for which there is therefore no practical means of fitting the model to data. To best understand this estimation procedure and its importance to the present article, however, we’ll begin by first taking a step backwards and considering the most commonly used phylogenetic model for continuously valued phenotypic trait data, and that is the Brownian motion model (Felsenstein 1973; Felsenstein 1985; Martins and Hansen 1997; Garland and Ives 2000; O’Meara et al. 2006; Harmon 2019).

Brownian motion is a continuous time directionless random walk or stochastic diffusion model that can be fully described by two parameters: *x*_0_, the starting value; and *σ*^2^, the instantaneous diffusion rate of the process (Felsenstein 1973; O’Meara et al. 2006; Revell and Harmon 2022). When applied to a single trait dimension, evolution via Brownian motion will tend to result in an increase or decrease of the trait with equal probability, variance along each lineage accumulates at rate *σ*^2^, and the variance after time *t* has a value of *σ*^2^ multiplied by *t* (O’Meara et al. 2006; Revell and Harmon 2022). Brownian motion is a Gaussian process, meaning that the distribution is normal with expectation *x*_0_ and variance *σ*^2^ *× t*, i.e., *∼ N* (*x*_0_*, σ*^2^*t*) (O’Meara et al. 2006; Revell 2008; Harmon 2019).

Figures 1a and 1b show (first) a simple phylogeny containing two terminal taxa (*T1* and *T2*) descended from a common ancestor, and (second) a single realization of unbounded Brownian motion evolution along the branches of this tree with a starting value of *x*_0_ = 0.0, and a diffusion rate parameter value of *σ*^2^ = 0.8.

**Figure 1:**
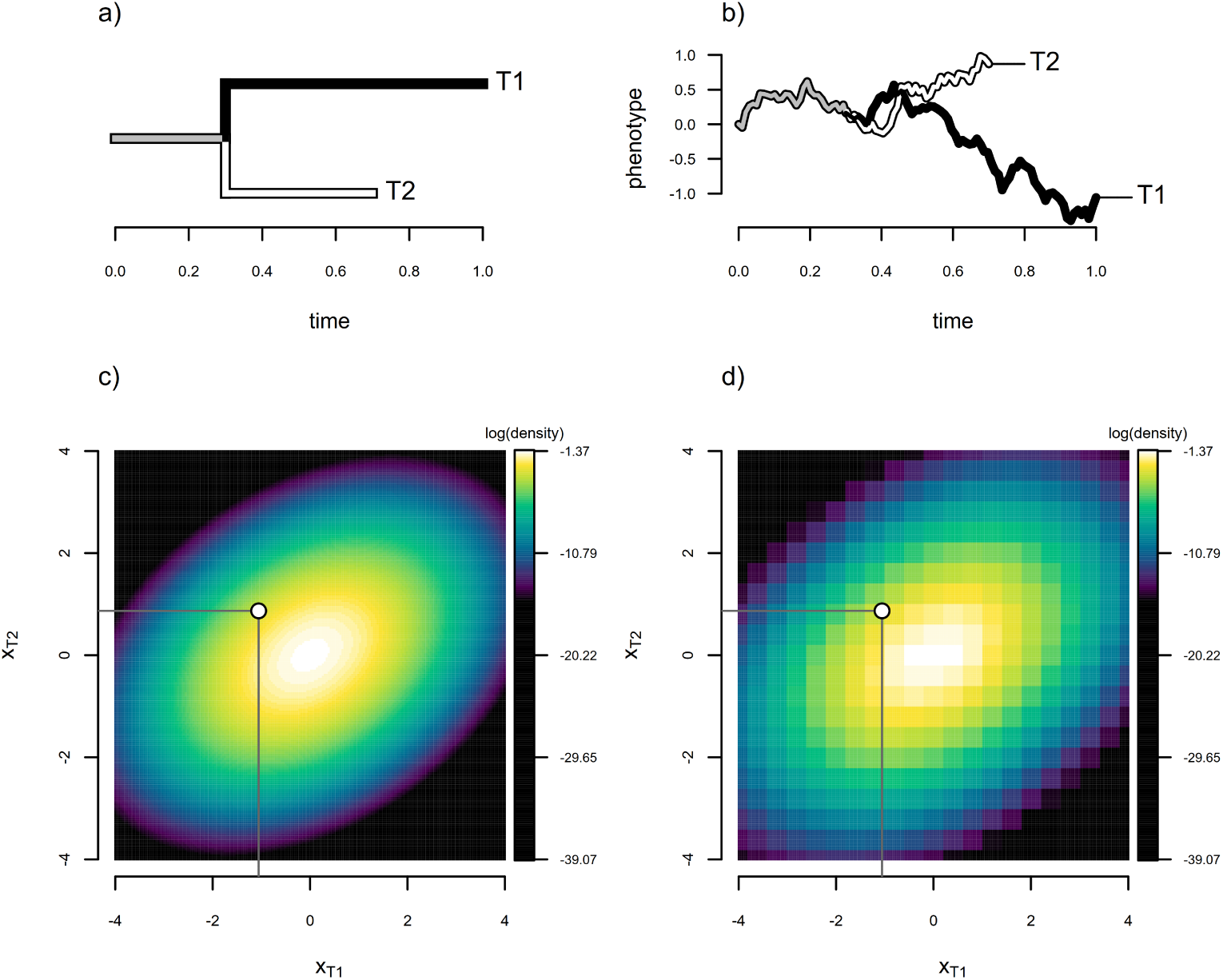
Illustration of the discretized diffusion or finite-state approximation of Boucher and Démery (2016) applied to a simple two-taxon case. a) Two-taxon phylogeny containing operational taxa *T1* and *T2*. b) Realization of unbounded Brownian motion in one dimension with starting value *x*_0_ = 0.0 and diffusion rate *σ*^2^ = 0.8. c) Exact probability density of the evolutionary process simulated in panel (b) with the simulated tip values, **x**, overlain, where *x*_0_ and *σ*^2^ have been set to their generating values. d) Density function obtained by discretizing the trait space and approximating the density from the probability of the discretized data under a continuous time ordered Markov chain on a grid, that is, using the discretized diffusion approximation. As for panel (c), here *x*_0_ and *σ*^2^ have been set to their generating values.

When evolution occurs via Brownian motion, as shown in Figure 1b, the probability density of species data on a phylogenetic tree is multivariate normal with variances equal to the total height of each taxon multiplied by *σ*^2^, and covariances between each pair of taxa (e.g., *T*_1_ and *T*_2_ in our simple case) equal to *σ*^2^ times the height above the root of their common ancestor (Freckleton et al. 2002; O’Meara et al. 2006; Revell 2008; Revell and Harmon 2022). That is to say, **x** *∼* MVN(**x**_0_*, σ*^2^**C**), where **x** is a vector of species values, **x**_0_ is a repeating vector of the root state, *x*_0_, **C** is an *N × N* matrix containing the height above the root of the common ancestor of each pair of taxa (in the off-diagonals, and the total height of the tree to each taxon in the diagonal), and *σ*^2^ is as defined previously (O’Meara et al. 2006; Revell 2008; Harmon 2019; Revell and Harmon 2022).

In the example of Figure 1a, this gives 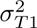 = 0.80, 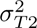 = 0.56, and a covariance between species equal to the amount of common ancestry they share multiplied by *σ*^2^, here *σ_T_* _1_*_,T_* _2_ = 0.32 (Figure 1a). To calculate the probability density of any **x** from *N* species given values for *σ*^2^ and *x*_0_, then, we merely evaluate the corresponding *N* dimensional multivariate normal distribution, which can be done on a computer using standard numerical methods for trees containing up to thousands of taxa, with fast methods available for even larger phylogenies (e.g., Ho and Ané 2014a). This calculation is illustrated on a natural logarithm scale in Figure 1c for the phylogeny and generating parameter values (*x*_0_ and *σ*^2^) of Figures 1a and 1b.

If the values of *x*_0_ and *σ*^2^ were not known, as would typically be the case in an empirical study, this is the same probability density function that we’d use if we aimed to identify the values of *x*_0_ and *σ*^2^ that made our observed data most probable: that is, for Maximum Likelihood estimation (Pagel 1999; Revell and Harmon 2022). In that case, we’d merely evaluate the probability density at *x_T_* _1_ and *x_T_* _2_ for different values of *σ*^2^ and *x*_0_ until we identified those that maximized the probability density of our observed data in **x** (Lynch and Walsh 1998; O’Meara et al. 2006; Revell and Harmon 2022). (In practice, of course, we have analytic solutions for the Maximum Likelihood values of *x*_0_ and *σ*^2^, e.g., see O’Meara et al. 2006, but the principle remains the same.)

Boucher and Démery (2016) appreciated that because a continuous time, discrete state, ordered and symmetric Markov chain (also known as a symmetric nearest-neighbor Markov chain, or a continuous-time random walk on a lattice) converges to Brownian motion in a single trait dimension in the limit as the number of levels in the Markov chain goes to *∞* (Donsker 1951; Billingsley 1999), one should be able to compute an approximation of the probability density of a Brownian process after first discretizing the continuous trait state space into *n* bins each of width *δ* (Boucher and Démery 2016; Boucher et al. 2018). This approximation will become exact as *n → ∞* (or, equivalently, as *δ →* 0). We refer to this model as discretized stochastic diffusion because it should behave in precisely the same way as Brownian motion in the limit as the step interval width of our trait space, *δ*, goes towards zero. (Technically, this exact equivalence also requires that our trait space go from *−∞* to *∞* for an unbounded Brownian process, although in practice it is very nearly exactly equivalent so long as our trait space is substantially larger than the probable trait range given *x*_0_, *σ*^2^, and the total time represented by our tree.)

Although the application of this approach in Boucher and Démery (2016) was, to our knowledge, novel in phylogenetic comparative methods, the discretized diffusion approximation can also be viewed as a finite-difference representation of the diffusion generator (closely related to finite-difference discretizations of the forward and backward Kolmogorov equations, see Boucher et al. 2018). As the discretization step *δ →* 0, this approximation converges to the underlying continuous diffusion process, a well-established result in the theory of stochastic processes (Ethier and Kurtz 1986; Risken 1996; Kushner and Dupuis 2001; Gardiner 2009; Pavliotis 2014; Boucher et al. 2018).

To exploit this convergence to calculate the probability density of our original continuous data (**x**), Boucher and Démery (2016) realized, one begins by computing the probability of the now discretized data (let’s refer to them as **x***^′^*) as if they’d arisen under a continuous time, ordered, Markov chain with a rate of change between adjacent levels equal to *q_c_*= *σ*^2^*/*(2*δ*^2^), where *δ* is (again) the bin width of our trait space discretization, and *σ*^2^ is our stochastic diffusion (in this case, Brownian motion) rate. This construction corresponds to a classical approximation in which a diffusion process is represented by a nearest-neighbor continuous-time Markov chain on a lattice, with the approximation converging to the continuous Brownian diffusion process as *δ →* 0 (e.g., Ethier and Kurtz 1986; Kushner and Dupuis 2001; Pavliotis 2014). The probability of the discretized data, **x***^′^*, can then be computed efficiently using the pruning algorithm of Felsenstein (1981, 2004; Yang 2006).

The probability *density* of the data, *f* (**x**) (as opposed to the *probability* of the data after discretization, *p*(**x***^′^*)), is then approximated as *f*^^^*_δ_* (**x**) = *p*(**x***^′^*)*/*(*δ^N^*) (where *f*^^^*_δ_* (**x**) *→ f* (**x**) as *δ →* 0), in which (again) **x***^′^* contains the discretized values of our continuous trait *x*, *p*(**x***^′^*) is the probability of **x***^′^* under the Markov chain model, *N* is the number of operational taxa in our tree, and *δ* is as it’s been defined previously. This relationship follows from the standard connection between probabilities of histogram bins and the corresponding continuous probability density (e.g., Silverman 1986).

An approximated probability density obtained using a discretization of our trait space into a mere *n* = 21 continuous character bins (far fewer than we’d employ if applying this in practice) for our example of Figure 1 is shown in Figure 1d. The resemblance of this surface to the analytic equivalent in Figure 1c is already quite apparent, even for this purposefully “rough” discretization. That is to say, it already seems clear that we might be able to approximate a continuous density function by substituting a finite-state discretization, in this case, a symmetric Markov chain on a grid, for the original continuous one, Brownian motion.

Figures 2a and 2c show the equivalent discretely approximated probability density function as Figure 1d, but where the number of levels of the discretization has been roughly doubled (to 41, and *δ* thus halved from its previous level) in panel 2a and doubled again (to 81) in panel 2c. Figures 2b and 2d give the difference from the analytic density of Figure 1c (Δ density) for each of the two preceding panels. Unsurprisingly, the accuracy of our approximation (i.e., its resemblance to Figure 1c) concomitantly improves as the number of bins is increased (in other words, as *δ* is decreased towards zero; Figures 2b, 2d).

**Figure 2:**
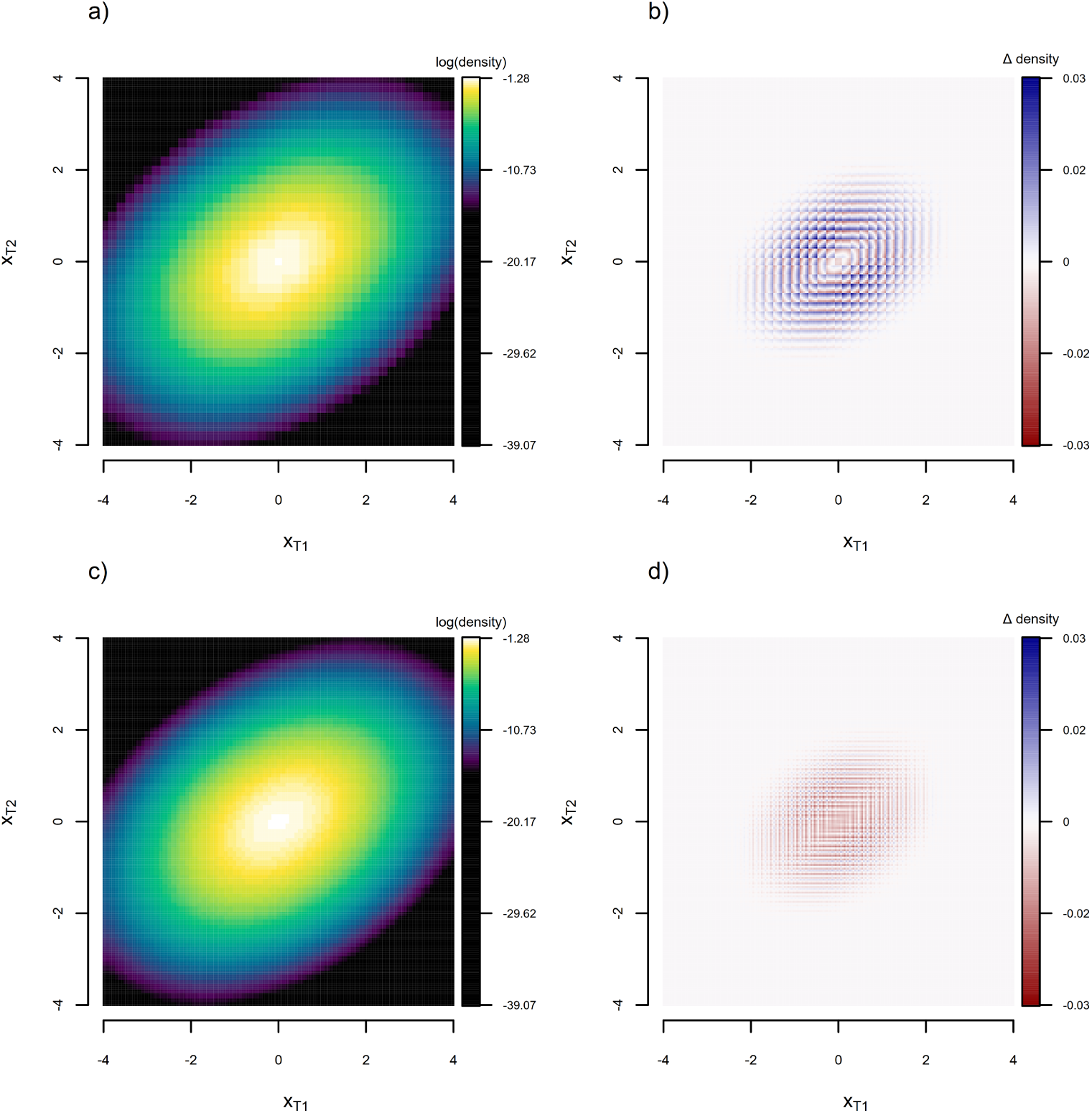
a) Density function obtained for the tree and generating stochastic process of Figure 1 using the discretized diffusion approximation with 41 levels, under the generating parameters of the process (*x*_0_ = 0.0 and *σ*^2^ = 0.8). b) Difference between the true (i.e., analytic) probability density of Figure 1c and the estimate obtained via the discretized diffusion approximation in panel (a). c) The same as (a), but with about twice as many bins. The increasing resemblance to Figure 1c is apparent. d) Difference between the true and estimated density for panel (c).

What we’ve illustrated here is simply that the probability density of Brownian motion on a tree can be approximated by discretizing the trait space, computing the probability of our now discrete continuous character data under a scenario of discretized symmetric stochastic diffusion, and then applying the appropriate back-transformation (from probability to density function), as previously documented (Boucher and Démery 2016; Boucher et al. 2018).

At this point we think it could be useful to highlight a common misconception about our approach. Frequently, colleagues misunderstand the preceding explanation of the discretized diffusion or finite-difference approximation as implying that we’ve proposed to transform a continuous problem into a discontinuous one: akin to recategorizing continuously varying lizards into “small,” “medium,” and “large,” and then proceeding to analyze these “new” data using machinery suitable for a discrete trait. This is wrong. What we’re describing instead is that it’s possible to exploit the equivalence (in the limit as *δ →* 0) of Brownian motion and an ordered, symmetric, continuous time Markov chain on a lattice to calculate an estimate of the probability density of our original, untransformed, continuous trait model – using an approximation that, once again, converges in the limit as *δ*, the interval width of our discretization of continuous trait space, shrinks towards zero.

This would be no more than an interesting curiosity (for those of us who work in phylogenetic comparative methods) if it could only be applied to trait evolution models, like unbounded Brownian motion, for which an analytic solution for the probability density is known and numerically tractable. To appreciate the proper utility of the Boucher and Démery (2016) approximation, therefore, we must identify and apply it to evolutionary scenarios where an analytic function giving the relationship between data and the probability density is unavailable or impractical to use, but for which a version of the discretized diffusion approximation can be constructed (Boucher et al. 2018).

Before we continue to consider the various such scenarios listed in the Abstract of this article, we’ll commence with the canonical case of bounded Brownian motion with reflective boundary conditions (Boucher and Démery 2016). Here, there’s no convenient analytic density (because the analytic solution of Boucher and Démery 2016 scales poorly to reasonably-sized phylogenies), and the underlying process is non-Gaussian due to the boundaries of trait space. This means that an analytic density, if it could be evaluated, would not be multivariate normal. Nonetheless, this evolutionary scenario poses no difficulty at all to our approximation because it’s straightforward to compute the probability of data arising under an ordered Markov chain with reflecting limits, and this is all that’s required to evaluate the likelihood using our discretized diffusion approximation (Karlin and Taylor 1981; Risken 1996; Boucher and Démery 2016). In practice, this is done merely by removing outward transitions at the bounds of the trait space. The missing outward rate can either be reassigned to the inward transition, resulting in a boundary rate of 2*q_c_* = *σ*^2^*/δ*^2^, or simply omitted, leaving the inward rate equal to *q_c_* = *σ*^2^*/*(2*δ*^2^) in the boundary rows. The former is more technically accurate for a finite number of bins of discretization, *n*, but the latter is computationally simpler and converges to the former as *δ →* 0. (Both approaches are implemented in the *phytools* R package of Revell (2024); however, we use the latter in all of our analyses for computational efficiency.)

Figure 3a gives a single realization of bounded Brownian evolution for *σ*^2^ = 3.0, *x*_0_ = 0, and bounds [*−*2, 2] on the same tree as was shown in Figure 1a. Figure 3b shows the probability density under a bounded Brownian process, given the generating parameter values of our simulation, and obtained using 101 bins of discretization. It’s easy to appreciate from the figure that this density is not bivariate normal, as we saw in Figures 1 and 2, but saddle-shaped (Figure 3b). Just as before, under the more typical conditions where the generating parameters of the stochastic process were unknown, we would just evaluate the density function at *x_T_* _1_ and *x_T_* _2_ for different values of *σ*^2^, *x*_0_, and bounds until we found those that made our observed data most probable. These would be our Maximum Likelihood estimates (Boucher and Démery 2016). (Note that, in practice, Boucher and Démery 2016 have shown the uppermost and lowermost observed values in **x** tend to fully determine the MLE bounds in this model: i.e., the ML estimate of the bounds will be the observed trait range, unless otherwise specified.)

**Figure 3:**
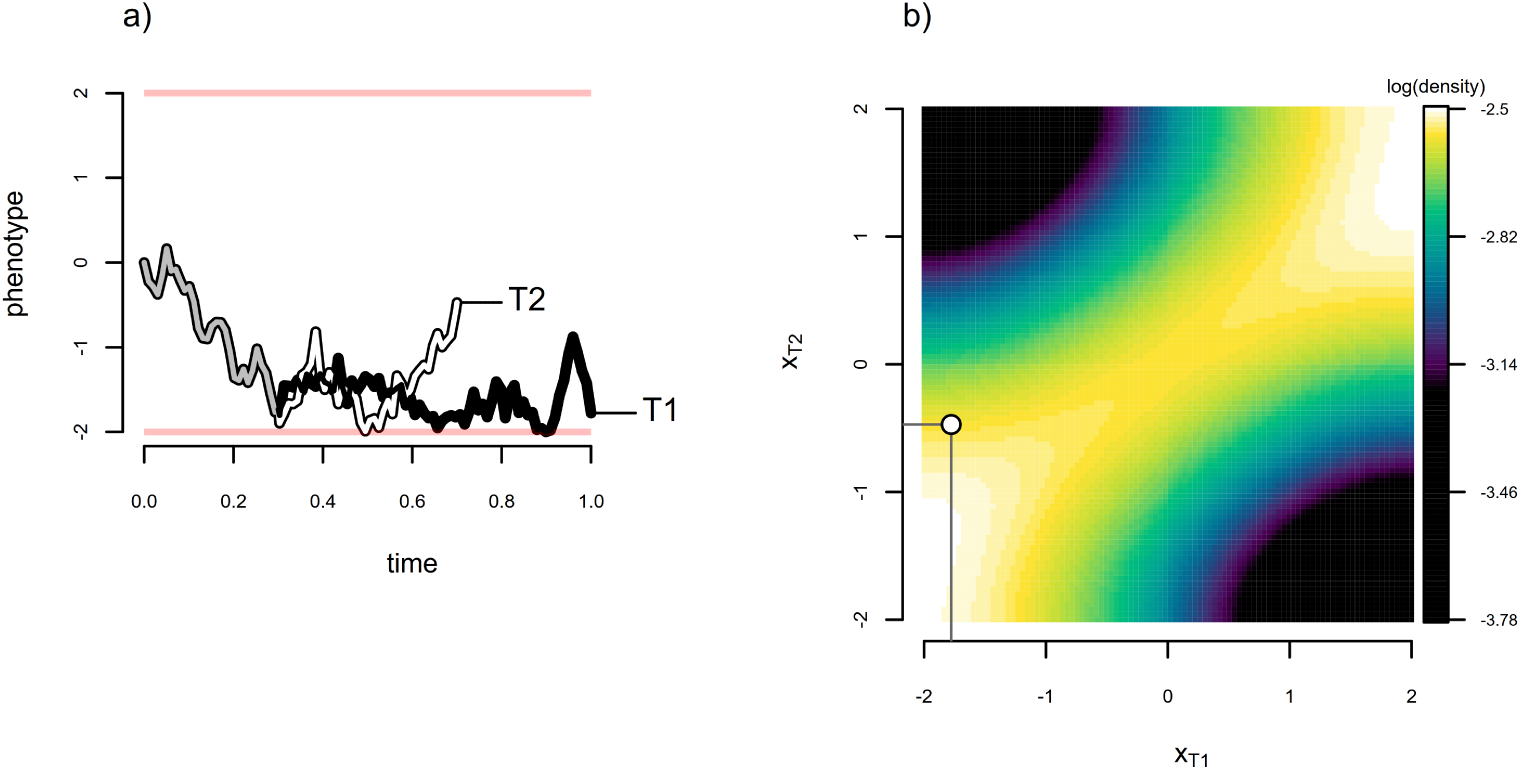
Demonstration of the application of the discretized diffusion approximation to a case (bounded Brownian motion) for which an analytic probability density is not readily available. a) Realization of Brownian motion evolution with reflective bounds on the phylogenetic tree of Figure 1a. Reflecting boundaries are shown by the red bars at the top and bottom of the phenotype space, [*−*2, 2]. b) Probability density function estimated using the discretized diffusion approximation under the generating parameter values of figure panel (a).

To demonstrate that our discretized diffusion approximation has provided us with an accurate measure of the probability density under our process, we undertook 2,000,000 simulations of bounded Brownian evolution on the two-taxon phylogeny of Figure 1a using the same conditions as were used to obtain the data of Figure 3a: i.e., *σ*^2^ = 3, *x*_0_ = 0, and reflecting bounds at [*−*2, 2]. We then plotted the raw simulation results using semi-transparent colors (in Figure 4a) and computed a two-dimensional kernel density, which we have graphed on a natural logarithm scale (Figure 4b). The similarity of this kernel density and our probability density function estimated using the discretized diffusion approximation in Figure 3b is striking.

**Figure 4:**
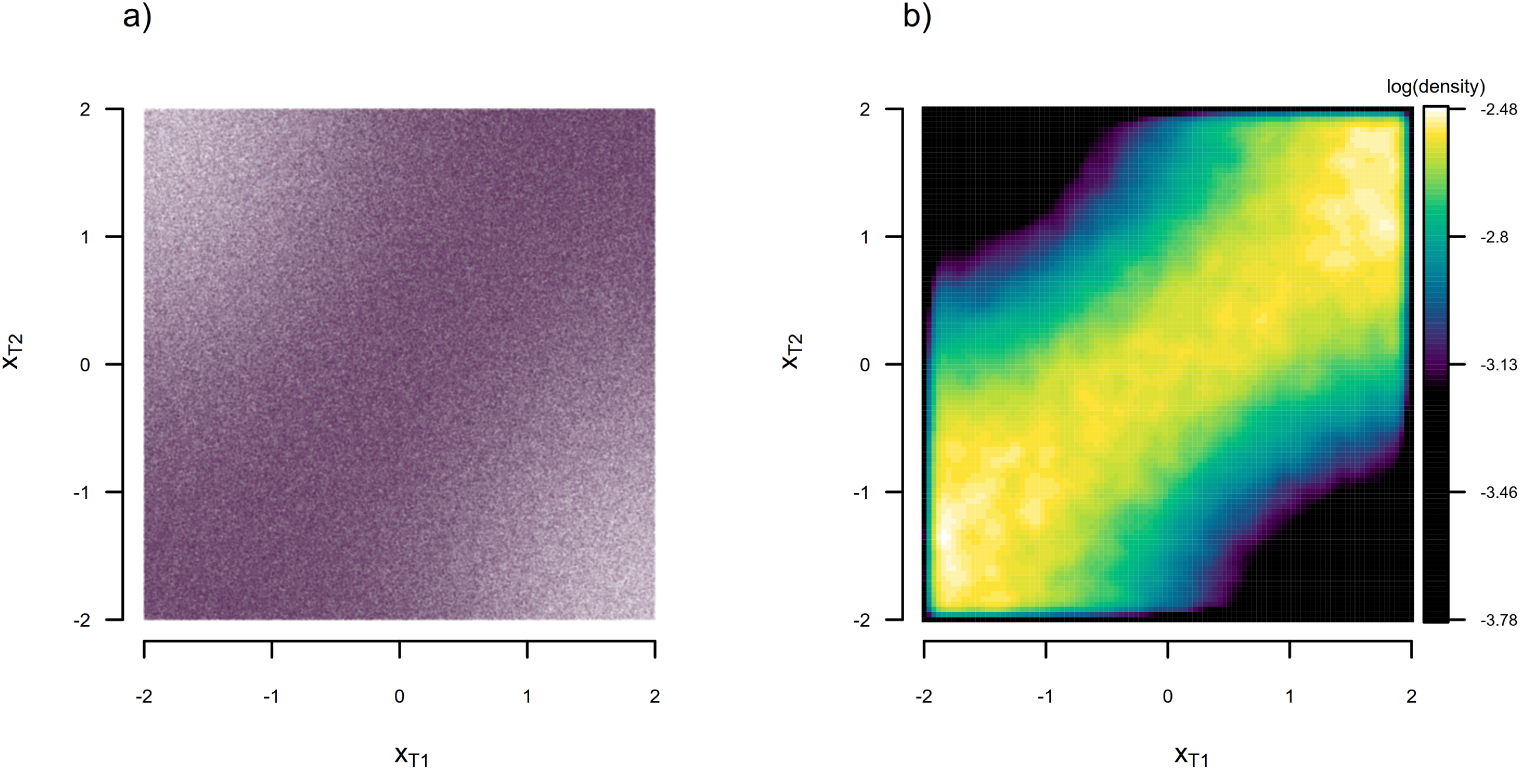
a) Data for 2,000,000 bounded Brownian motion simulations on the phylogeny of Figure 1 using the simulation conditions of Figure 3a. b) Two dimensional kernel density visualization of the data of panel (a). Note the resemblance of this kernel density to the probability density surface obtained via the discretized diffusion approximation and illustrated in Figure 3b.

Finally, to illustrate that the discretized diffusion approximation is also useful in estimation (that is, in fitting this model to data), we undertook a very small simulation analysis. We began by simulating one hundred 200-taxon stochastic pure-birth phylogenetic trees. We then randomly sampled generating values of log(*σ*^2^) for bounded Brownian evolution from a normal distribution with mean of zero and variance of one, and simulated continuous trait data for each tree using reflecting bounds of [*−*1, 1]. Then we proceeded to fit both standard (unbounded) Brownian motion, using a standard analytic function for the likelihood, and Boucher and Démery (2016)’s bounded model, using the discretized diffusion approximation exactly as described above. Figure 5 shows parameter estimation for *σ*^2^ and *x*_0_ for each of these analyses. (For a more comprehensive set of simulations and analyses under this model, we refer the reader to Boucher and Démery 2016.)

**Figure 5:**
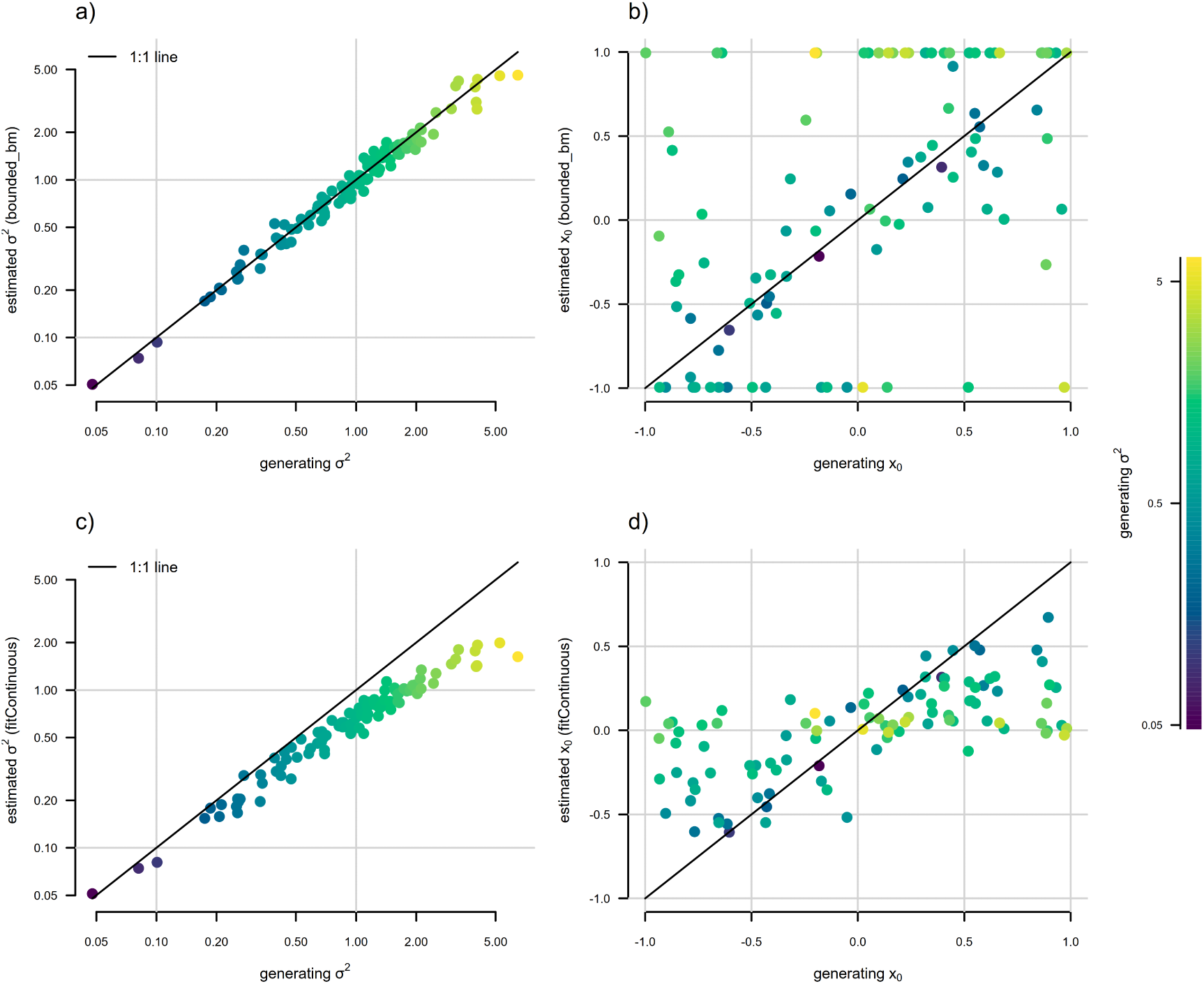
Statistical behavior of Boucher and Démery’s (2016) bounded Brownian motion model, fit using the discretized diffusion approximation. The point color gradient corresponds to the generating stochastic diffusion rate, *σ*^2^, with higher values of *σ*^2^ shown by lighter colors, as illustrated in the right figure legend. a) Generating vs. estimated *σ*^2^ obtained by fitting a bounded model following Boucher and Démery (2016). b) Generating vs. estimated *x*_0_ from the same bounded model as in panel (a). c) Generating vs. estimated *σ*^2^ when standard (unbounded) Brownian evolution was assumed. d) Generating vs. estimated *x*_0_ under unbounded Brownian motion. Bounded Brownian motion parameter estimates were obtained using the *phytools* R package (Revell 2024), while unbounded estimates come from the *geiger* package (Pennell et al. 2014). See main text for more details.

In general, we found that when trait data were simulated under bounded Brownian motion, ignoring the bounds in model-fitting resulted in systematic underestimation of *σ*^2^, particularly for larger values of the rate parameter (Figure 5c). This downward bias arises because the reflective bounds decouple the diffusion rate, *σ*^2^, from the amount of phenotypic dispersion that can accumulate under the process. The bias completely disappears when we instead fit Boucher and Démery (2016)’s model using the discretized diffusion approximation (Figure 5a).

Both bounded and unbounded Brownian motion resulted in reliable estimation of the root state, *x*_0_, for low values of *σ*^2^ (dark colored points in Figures 5b and 5d); however, point estimates of *x*_0_ tended towards the upper or lower bound when the bounded model was used for estimation, and towards the midpoint of the bounds (i.e., zero for the evolutionary scenario of our simulations) when unbounded Brownian evolution was assumed (light colored points in Figures 5b and 5d). Nonetheless, prior work has shown that confidence intervals for the bounded model are correct and estimation error tends to be quite a bit smaller when the data arose via a bounded process (Revell 2025).

Now that we’ve reviewed this canonical model and re-established that the discretized diffusion approximation estimation procedure of Boucher and Démery (2016) is useful for the bounded case, we’ll proceed to consider a small number of other applications, as described in the Abstract.

## 2 Applications

Having now described the basis of the discretized diffusion approximation as an estimation method for models without closed form or tractable solutions for the likelihood, and having established its usefulness for fitting a bounded Brownian motion process to phylogenetic data, we now continue to apply it to several models for discrete, continuous, or jointly evolving discrete and continuous traits. Of these, at least four are partly or wholly new to phylogenetic comparative biology. To reiterate a point made earlier, but one that bears repeating due to its importance, in each case discretization is being used as a numerical approximation to the true underlying probability distribution or probability density under the original evolutionary process, not as a simple recoding of continuous observations into categorical data.

For each evolutionary process, we commence by describing the model as a hypothetical generating process of real biological data. We then proceed to characterize the finite-state representation of this model that we propose to use for estimation. We continue to use numerical simulations to either verify convergence with an independently calculated likelihood (where available), to examine model parameter estimation, or to compare the method to existing estimation procedures. In line with the purpose of this article, these analyses are intended as a proof of concept for each model, rather than as a comprehensive investigation of statistical properties or performance. We intend to subject each of these models to more thorough study in subsequent research. In the following subsections we’ll consider the threshold model, a new semi-threshold model, and then three different joint discrete and continuous trait models.

### 2.1 The threshold model

The threshold model gives a scenario in evolutionary quantitative genetics positing that although our trait may be observed only in discrete conditions (e.g., 0 or 1; white, red, or blue; present or absent; etc.), its expressed value is based on some underlying, unobserved continuous character called liability (Wright 1934; Felsenstein 2005, 2012; Revell 2014; Revell and Harmon 2022). Liability is the unobserved attribute that actually evolves over the branches and nodes of our phylogeny, even though what’s observed at the tips of the tree is the discontinuous byproduct of this evolution (Felsenstein 2005; Revell 2014; Revell and Harmon 2022). Every time the unobserved trait liability crosses a fixed threshold value, the observed discrete character changes in state.

A well-established model in evolutionary quantitative genetics (Lynch and Walsh 1998), the threshold model was originally described to characterize the genetic basis for variation in digit number of guinea pigs (and other discrete traits with a similar hypothesized genetic basis) by Wright (1934), but was later introduced to phylogenetic comparative analysis by Felsenstein (2005, 2012; Revell 2014). A threshold evolutionary process might be expected for discrete characters whose underlying basis is determined by a potentially observable, but unmeasured, quantitative characteristic, like the level of a circulating blood hormone that triggers a binary developmental switch; but the threshold model could also capture the evolutionary dynamics of complex ecological characteristics, such as preferred habitat, that manifest or are measured discretely, but are actually underlain by many continuous and discretely-valued phenotypic attributes (Revell 2014; Revell and Harmon 2022).

An illustration of evolution under the threshold model is given for a simple three-taxon tree in Figure 6. Figure 6a shows the (normally unobserved) evolutionary trajectory of liability, whereas Figure 6b shows the resultant discrete character history of our observed trait.

**Figure 6:**
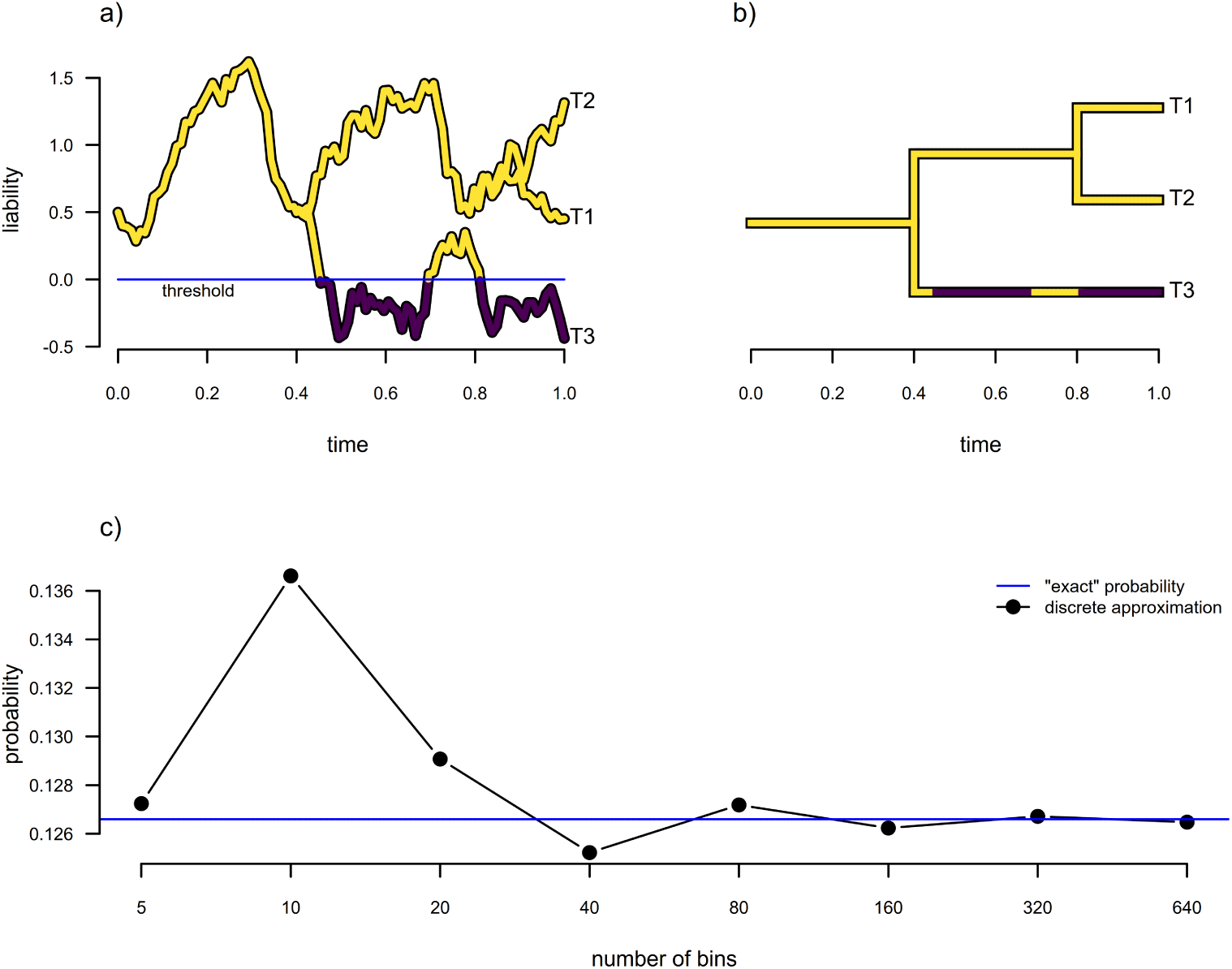
Estimation under the threshold model from evolutionary quantitative genetics. a) Evolution of the trait liability via Brownian motion. When liability crosses the “threshold” value of 0 (in this example), the discrete character changes in state. b) Corresponding discrete character phylogenetic history for the liabilities of panel (a). c) Comparison of “exact” probability of the discrete character data at the tips of the tree in panel (b) to the discrete approximation for various numbers of bins. See main text for more details.

Computing the probability of data that arise under a threshold model on the phylogeny normally requires that we calculate the integral of an *N* dimensional multivariate normal distribution (for *N* terminal taxa in our tree), which can be done efficiently using numerical methods for only relatively low *N* (e.g., Genz 1992, 2004). More efficient algorithms for computing higher dimensional multivariate normal integrals have been developed, but they still remain computationally impractical for trees containing the larger number of operational taxa that typify most modern phylogenetic comparative studies (Felsenstein 2005, 2012; Genz and Bretz 2009; Revell 2014).

In the original article describing application of the threshold model to phylogenetic comparative biology Felsenstein (2005) remarked that to compute the probability of the observed (discrete data) we must approximate “a high-dimensional integral for which no closed-form formula exists. . . . [t]he integral is approximated by Monte Carlo integration – drawing a large sample of points from the domain of integration and evaluating the contribution to the integral in the vicinity of these points by evaluating the function at the points. we replace the continuous function that is being integrated by a histogram that approximates it.” Felsenstein (2005) proceeds to describe how this can be done efficiently using Markov chain Monte Carlo with Gibbs sampling (McCulloch 1994; Felsenstein 2005).

Based on Boucher and Démery (2016), it occurred to us that this integral could also be approximated in a different way — by replacing our underlying continuous stochastic process for liability with a discretized diffusion approximation that converges on the continuous one as our step size shrinks, *δ →* 0, just as we’ve described above. In Figure 6c the equivalence of our “exact” solution (obtained using numerical integration for the very small *N* = 3 tree of Figure 6b) and the discrete approximation in the limit as the bin number is increased (and thus their widths, *δ*, decreased towards zero) is apparent.

To illustrate the usefulness of this approach for fitting a multi-state threshold model to comparative data, we undertook a small simulation study. We first generated one hundred 200-taxon stochastic pure-birth phylogenetic trees, each scaled to 3.0 units of total depth. We next simulated liability evolution on each tree under Brownian motion with *x*_0_ sampled randomly on the interval [*−*1, 3] and *σ*^2^ = 1.0. We created a discrete threshold character with up to four states from these simulated liability values using the thresholds of [0, 0.8, 2] between the discrete character levels *a* and *b*, *b* and *c*, and *c* and *d*, respectively, although in some cases not all four character states were observed among living taxa of our simulation. We proceeded to fit a multi-state threshold model (using the discretized diffusion approximation), as well as both symmetric and asymmetric ordered M*k* models (the two flavors of M*k* model that we felt most closely resembled an ordered multi-state threshold process), to each simulated tree and dataset (Pagel 1994; Lewis 2001; Beaulieu et al. 2013; Revell and Harmon 2022).

As a side note, the multi-state threshold model is only identifiable if we fix *σ*^2^ and one of the thresholds (Felsenstein 2005; Revell and Harmon 2022). This is because the liabilities are unobserved and therefore have no natural units or starting point. Absent these constraints, the liability scale, threshold positions, and diffusion rate could all be shifted or rescaled together without changing the probability of the observed discrete tip states. Following convention, we fixed *σ*^2^, the rate of evolution of the liability, to *σ*^2^ = 1, and the threshold between two adjacent character levels (in this case, *a* and *b*) to be 0. Doing so allows the remaining thresholds and root state to be estimated relative to these quantities (Revell 2014; Revell and Harmon 2022).

From the results of each simulation and analysis we first extracted the Akaike model weights of each of our three fitted models, and then the Maximum Likelihood estimates of the relative positions of the remaining two estimable thresholds between discrete character levels. Results from this analysis are shown in Figure 7.

**Figure 7:**
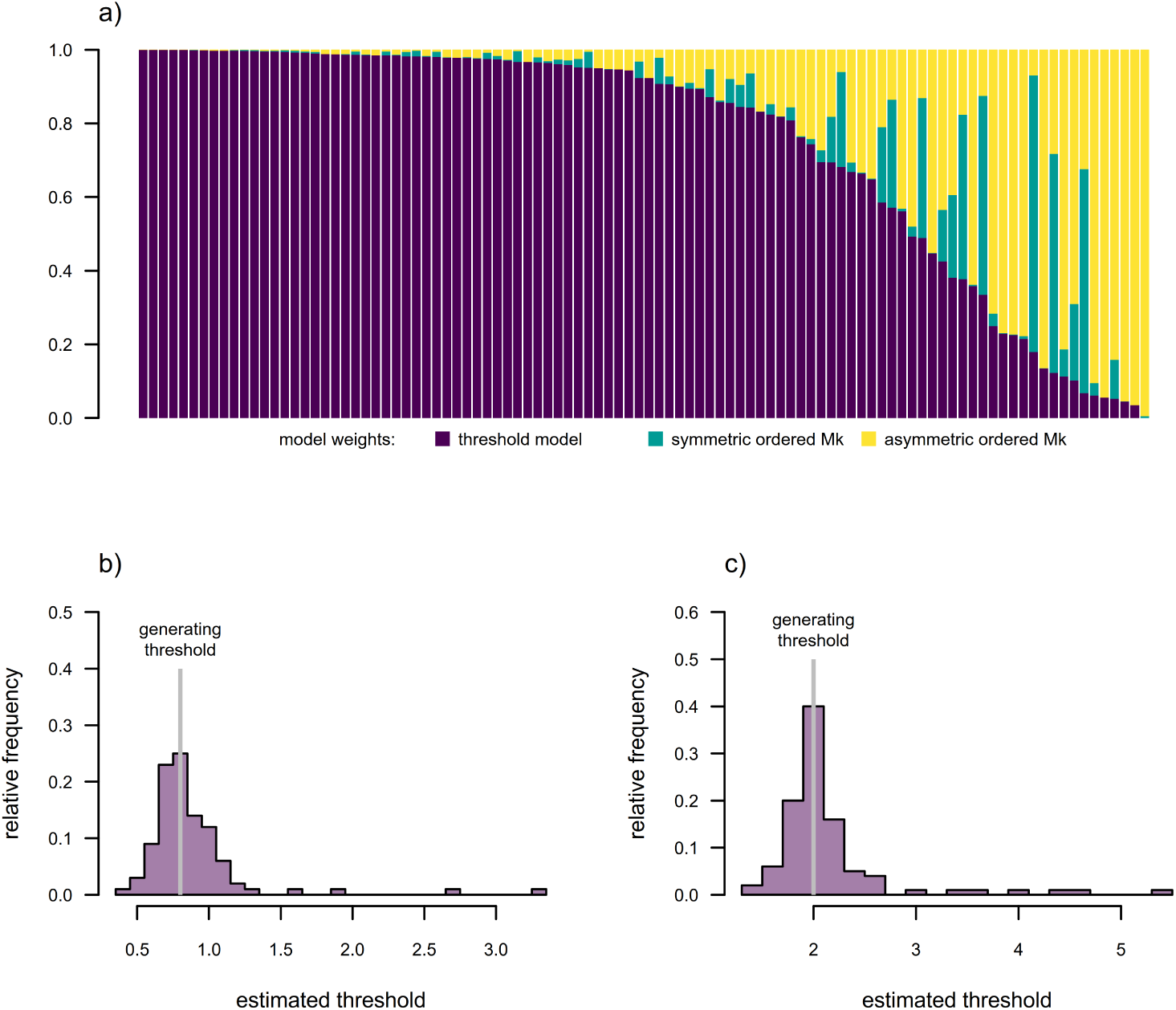
Results for model selection and parameter estimation for the multi-state threshold model using the discretized diffusion approximation for model-fitting. a) Akaike model weights for the threshold model and two variants of the M*k* model: an ordered symmetric model; and an ordered asymmetric model (the two flavors of M*k* model we thought most-closely resembled an ordered threshold process). Simulation model fits have been sorted by the model weight favoring the threshold model, which was the generating model for all simulations. b) Distribution across simulations of the estimated position of the threshold between character levels *b* and *c*, where the generating threshold was 0.8. (The position of the threshold between *a* and *b* was fixed to 0.) c) Distribution across simulations of the estimated position of the threshold between character levels *c* and *d*, where the generating threshold was 2. All threshold model parameter estimates were obtained using the *phytools* R package (Revell 2024) under the discretized diffusion approximation. See main text for additional details.

In general, we found that when data were simulated under a threshold model, the same model had the highest Akaike weight in 78% of simulations, and had at least 0.75 model weight in nearly ^2^ of all analyses (Figure 7a). We also found that the distribution of estimated thresholds tended to be centered^3^on the generating values of the two estimable thresholds in the model (Figures 7b, 7c). (This is not shown, but smaller experiments also indicate that for discrete data simulated under an ordered symmetric or asymmetric M*k* process, exactly the inverse pattern of model selection would hold.) In sum, our simulation results show that the discretized diffusion approximation can be used to fit a multi-state threshold model to discrete character data, and that this model can be discriminated from alternative generating processes such as ordered M*k* models (Figure 7).

### 2.2 A new semi-threshold model

As we reviewed in the prior section, a threshold model is one in which the observed trait is discretely-valued but underlain by an unobserved continuous character referred to as liability (Wright 1934; Felsenstein 2005, 2012; Revell 2014; Revell and Harmon 2022). Here we define a “semi-threshold” model, on the other hand, as one where we imagine that, on some part of its range, liability can be directly observed as a measurable quantitative trait. After crossing a given threshold, however, the character is fixed (while liability remains beyond the corresponding threshold) and liability itself can no longer be observed. Just as for the standard threshold model, trait liability, though no longer observable, nonetheless continues to evolve via the same stochastic process as on its observable range. As far as we know, this model is totally new to phylogenetic comparative biology. (At least one of us, LR, had previously discussed this model with J. Felsenstein in hypothetical terms; but we are not aware of any article describing it or any implementation of a model-fitting approach for this model in software.)

A simple example of a trait whose evolutionary dynamics could conceivably resemble this scenario might be “horn length” which can increase or decrease – but when it shrinks to zero, the horn is simply absent and can no longer be measured. The ability to produce a horn (now our unseen trait “liability”) might continue to evolve even if the horn is no longer present and measurable. (A species characteristic measured as the fraction of a whole, a phenotype that can only be observed in continuous form between 0 and 1 but that may nonetheless continue to change unobservably outside this range, is another example of a trait that could conceivably evolve via semi-threshold dynamics.)

Evolution under a semi-threshold process is illustrated by Figures 8a and 8b. Figure 8a shows the evolution of a single quantitative trait on the four-taxon tree of Figure 8b under a scenario in which the continuous character is observable on the interval [0, 1], but measurable only as being in condition 0 (or below, shown in black on the figure) or condition 1 (or above, shown in white) when outside this narrow range of values. The observable character history is projected onto the edges of the tree in Figure 8b.

**Figure 8:**
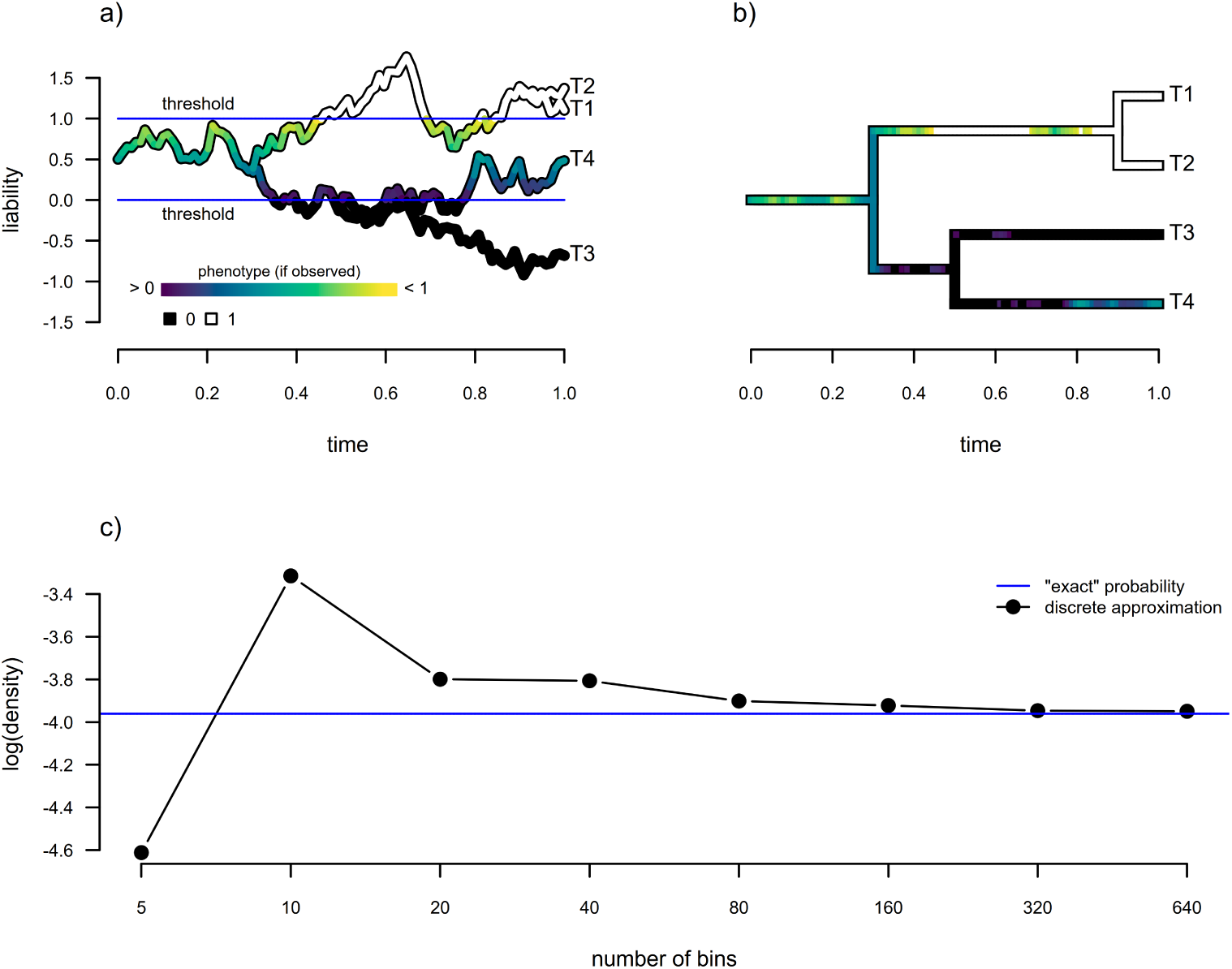
Estimation under a new semi-threshold model. a) Evolution of the quantitative trait or trait liability via Brownian motion. When the quantitative trait reaches either threshold of 0 or 1 (in this example), it becomes “stuck” on this value, even while liability continues to evolve. b) Corresponding continuous and discrete character phylogenetic history based on the trait evolution of panel (a). c) Comparison of “exact” log(probability density) of the continuous and discrete character data at the tips of the tree in panel (b) to the discrete approximation for various numbers of bins. See main text for additional details.

To compute the probability density of a set of data that we hypothesize have arisen under the semi-threshold scenario described above, we must evaluate an *N* dimensional multivariate normal density (for *N* taxa) over a region defined jointly by the observed (that is, non-thresholded) and censored (thresholded) values. For each tip whose trait falls within the observable range we evaluate the density at its observed value, whereas for tips falling below or above the threshold(s) we integrate the density over the corresponding semi-infinite interval (Tobin 1958). The likelihood is then given by the integral of the full multivariate normal over this mixed region of exact and truncated observations.

Fitting this model to data suffers from precisely the same difficulty as was described by Felsenstein (2005) for the standard threshold model in that no closed form solution exists for the integral of our multivariate normal density, and numerical methods to approximate it become unreliable for high *N*. Our new model, however, poses no difficulty whatsoever for the discretized diffusion approximation, with one small difference in implementation compared to the Brownian and bounded models previously illustrated. Specifically, if *m ≤ N* of the terminal taxa are represented by exact continuous observations, and the remaining *N − m* by censored values, the corresponding approximation to the mixed probability density must be obtained by dividing the probability of the discretized data, *p*(**x***^′^*), by *δ^m^* instead of by *δ^N^*.

Figure 8c illustrates the increasing resemblance between a log(density) computed from the discrete approximation, and the exact value obtained via numerical integration, as the number of bins is increased (i.e., for decreasing bin widths, *δ*).

To demonstrate parameter estimation under this new model, we undertook a small simulation study, similar to what we did for both the bounded model of Boucher and Démery (2016) and the multi-state threshold model from the prior section. In brief, we simulated one hundred 200-taxon stochastic pure-birth phylogenetic trees. For each simulated phylogeny, we randomly sampled a generating value of log(*σ*^2^) from a normal distribution with mean of zero and variance of one. We randomly sampled *x*_0_, the root state, uniformly on the interval [*−*1, 1]. Then, we simulated data under the semi-threshold model with upper and lower thresholds of [*−*1, 1]: meaning that the value of liability was observable on that interval, but simply measured as *−*1 or 1 outside of it. We proceeded to fit both the semi-threshold model, using our discretized diffusion approximation to compute the likelihood, or (for the purposes of comparison) a standard Brownian motion model to each simulated tree and dataset. Our results are shown in Figure 9.

**Figure 9:**
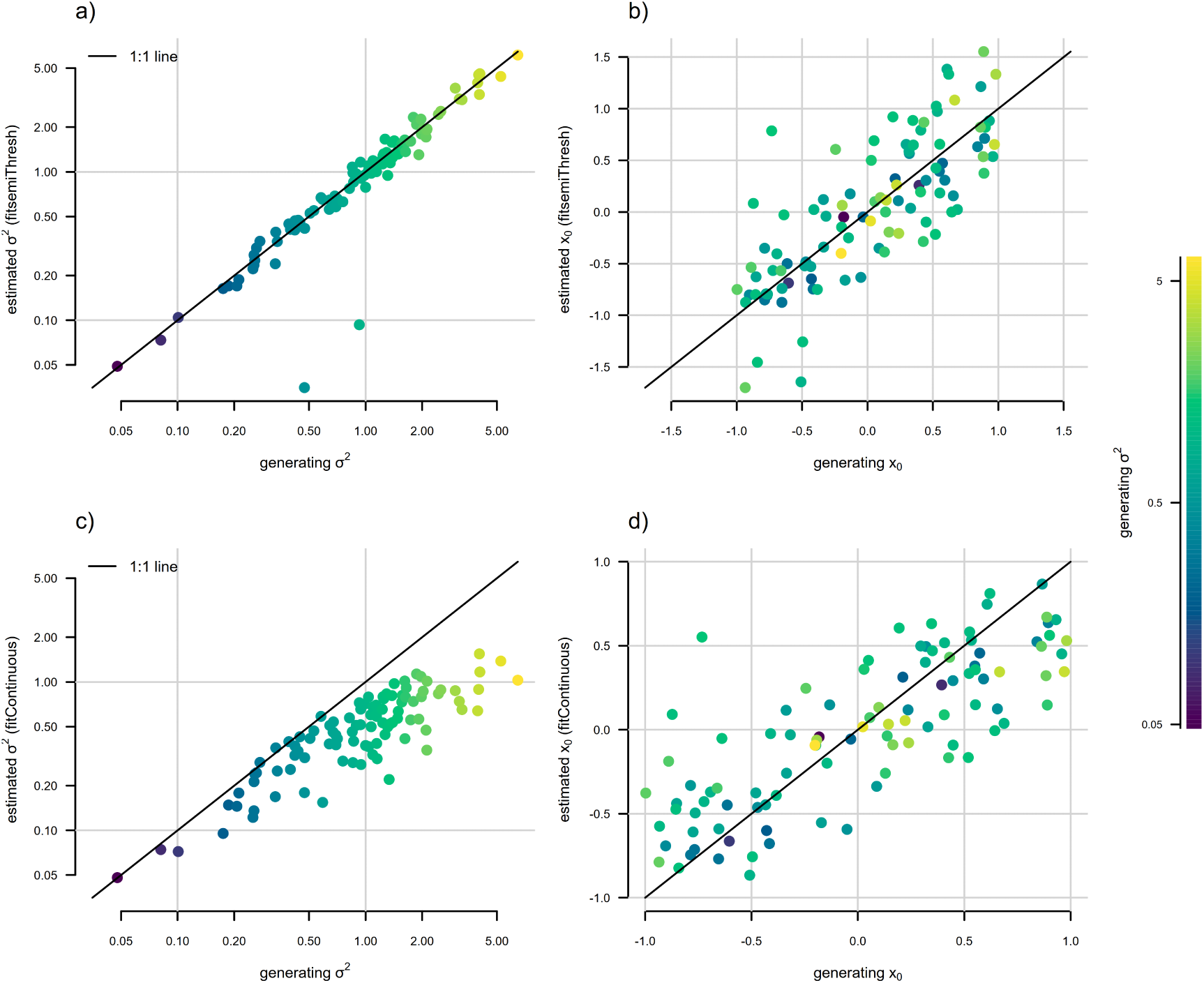
Parameter estimation under a new “semi-threshold” model for continuous trait evolution using the discretized diffusion approximation. The point color gradient corresponds to the generating stochastic diffusion rate, *σ*^2^, in which lighter colors match higher values of *σ*^2^. a) Generating vs. estimated *σ*^2^ obtained by fitting our new semi-threshold model using the discretized diffusion approximation. b) Generating vs. estimated *x*_0_ from the same semi-threshold model as in panel (a). c) Generating vs. estimated *σ*^2^ when standard Brownian motion evolution is assumed. d) Generating vs. estimated *x*_0_ under standard Brownian motion. Semi-threshold model parameter estimates were obtained using the *phytools* R package (Revell 2024) using the discretized diffusion approximation, while unbounded Brownian motion estimates come from the *geiger* package (Pennell et al. 2014), as in Figure 5. See main text for additional details.

In general, we found fairly reliable parameter estimation of both *σ*^2^ and (to a slightly lesser extent) *x*_0_ when the semi-threshold model was used for estimation (Figures 9a and 9b). By contrast, estimates of *σ*^2^ tended to be much less reliable under standard Brownian motion, and were biased downwards for increased *σ*^2^ (Figure 9c), just as we saw for the bounded model in Figure 5. Note that unlike the standard threshold model, *σ*^2^ appears to be estimable under this model, at least for the simulation conditions we explored, and we found no evidence of non-identifiability (Figure 9). We did not explore estimation of the threshold positions, since we assumed these would be known by the investigator (e.g., [0, 1] or [0*, ∞*], for a one-sided threshold). In some empirical circumstances in which a semi-threshold model was hypothesized, we can imagine the investigator setting one or the other bound to the limit of observed values of a trait. We’re uncertain of the consequences of so doing and would thus recommend that this be explored via further study.

In summary, in this subsection we have described a new trait evolution model for phylogenetic comparative biology called the “semi-threshold” model for tip data consisting of a mixture of continuous and censored observations. Much like the threshold model, computing the likelihood under this model is inherently challenging because it consists of integrating high-dimensional multivariate normal distributions. We show that the discretized diffusion approximation can be used to compute the likelihood under this evolutionary scenario, and when this approach is used on simulated data, our results show reasonable statistical properties in estimation (Figure 9).

### 2.3 A discrete-character-dependent continuous trait evolution model

A third model for which we realized that the discretized diffusion approximation could be put to use was the joint evolution of a discrete and a continuous trait, where the condition of the former affects the Brownian motion rate of evolution of the latter. Precisely this scenario is illustrated by Figure 10. Here, Figure 10a gives a discrete character history of a binary trait with two levels, *a* and *b*, evolving under a continuous time Markov chain (i.e., M*k* process, Lewis 2001); while Figure 10b shows the history of Brownian motion evolution of a continuous trait where 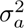 (the rate of continuous trait evolution when the discrete state is in condition *a*) is 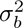 = 1.0, but 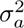 = 20.

**Figure 10:**
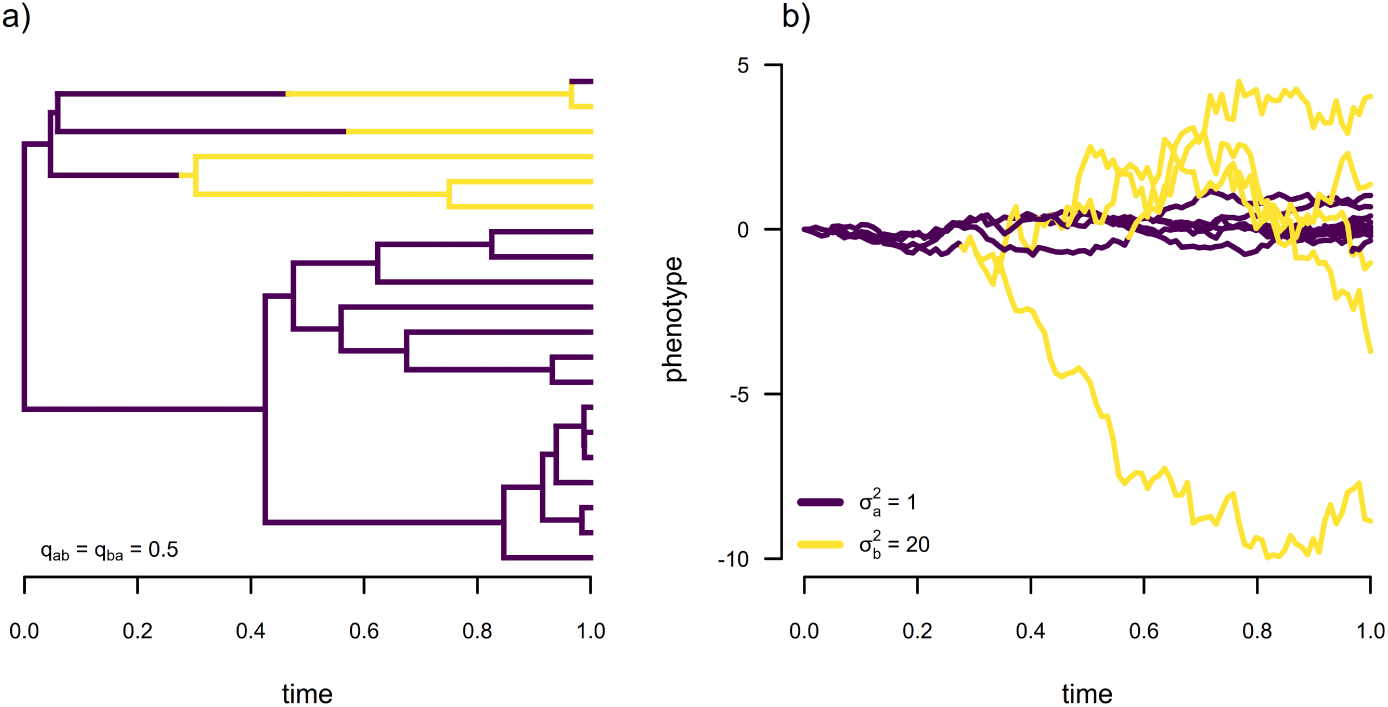
Discrete state dependent continuous character evolution. a) Simulated phylogenetic tree showing a discrete character history of a binary trait with two levels, *a* and *b*. b) Evolution of a continuous trait on the phylogeny of panel (a), where the rate of stochastic diffusion of the trait (*σ*^2^) varies as a function of the discrete trait such that 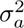 = 1.0 and 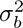 = 20.

A simple example of this sort of hypothesis might be one where different species of fish utilize pelagic (open water) or reef environments, and we’re interested in asking whether the rate of body shape evolution differs between these two sorts of habitat types (Friedman et al. 2020). Perhaps we expect much less shape evolution to occur per unit time in pelagic compared to reef habitats, due to the hydrodynamic constraints of fast open-water swimming (e.g., Friedman et al. 2020). Similarly, among polygynous and monogamous social mating systems in songbirds, we could hypothesize a higher rate of birdsong evolution in association with the former compared to the latter social mating system characteristic, due to stronger sexual selection on polygynous compared to socially monogamous male birds (e.g., Snyder and Creanza 2019).

The classic workflow to test hypotheses based on the evolutionary scenario of Figure 10 (or our hypotheses of habitat-dependent fish body shape evolution, or mating system dependent birdsong evolution, described in the previous paragraph), is a two-step procedure that involves first sampling a set of stochastic character mapped trees of our discrete trait in proportion to their probability following Huelsenbeck et al. (2003; Bollback 2006), fitting a rate-varying model to each sampled map and the continuous trait using the method of O’Meara et al. (2006; also see Thomas et al. 2006; Revell 2008; Revell and Harmon 2008), and then averaging across maps (e.g., see Revell and Harmon 2022). This workflow, though quite common in the literature, suffers from the problem that it will tend to underestimate the difference in continuous character rates between character levels of the discrete trait, particularly for high transition rates of our discrete trait (Revell 2013). We also suspect that it leads to less accurate estimation of the transition rates between character levels in the discrete trait itself, since the continuous character should also contain information about this process, but that information has been ignored in the two-step methodology.

At least two other notable related approaches have been proposed to address this problem. Firstly, May and Moore (2020) provide a Bayesian method to coestimate the transition process of a discrete character and discrete-state-dependent continuous character evolution with background rate variation. This method uses data augmentation of the discrete character data with unobserved discrete character histories over the tree and calculates an augmented likelihood as the product of the probability of the discrete character data and augmented history, times the conditional probability of the continuous trait data given the augmented history (May and Moore 2020). Similarly, Boyko et al. (2023) proposed a solution for jointly modeling discrete and continuous trait evolution that approximates the joint likelihood by adaptively sampling high-probability regime histories, informed by both the discrete and continuous traits in the model. Their approach does not allow for non-Gaussian continuous trait evolution processes and requires stochastic mapping as an internal step (Boyko et al. 2023). This might also be viewed as a different sort of approximation and is definitely not exactly equivalent to what we propose herein.

Our approach differs in that it involves calculating the joint likelihood of our discrete and continuous trait evolutionary model directly, using an approximation (the discretized diffusion approximation) that should become exact in the limit as the interval width of our discretization, *δ*, goes towards zero. Likewise, our model does not necessarily assume a Gaussian process of evolution for the continuous trait. It would be no trouble at all, for example, to fit this model to data assuming that evolution of the continuous trait occurred within reflective bounds, as in Boucher and Démery (2016), or under a semi-threshold process, like that described in the preceding section, among other evolutionary scenarios (e.g., Boucher et al. 2018).

To compute the probability under this model using the discretized diffusion approximation, we begin by building a sparse *k · n × k · n* transition matrix (referred to as an infinitesimal generator), for *k* levels of our discrete trait and *n* bins of discretization. Each of the *k · n* rows or columns represents a unique combination of a continuous character bin and discrete character level. Within each of *k* diagonal *n × n* blocks, the main super-and sub-diagonals of the generator are populated by the values 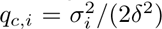 for each *i* through *k* discrete-state-dependent stochastic diffusion rates, 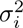. The diagonal entries of each non-diagonal *n × n* sub-matrix block, by contrast, contain the discrete character transition rates between each of the *k* levels of our discrete character. The main diagonals of the full matrix contain the negative row sums, and all other entries in the matrix are zero.

An illustration of this model structure is given in Figure 11 for 40 bins of discretization, although for accurate model parameter estimation a much finer discretization is recommended. To reiterate, in Figure 11, the macro-structure of this transition matrix consists of *k × k* = 4 quadrants: two (those along the diagonal) describing the evolution of the continuous character; and two (off the diagonal) describing the transition process for the discrete trait. As previously noted, each row and column of the matrix corresponds with one particular combination of a continuous character bin of width *δ* and discrete character state (Figure 11).

**Figure 11:**
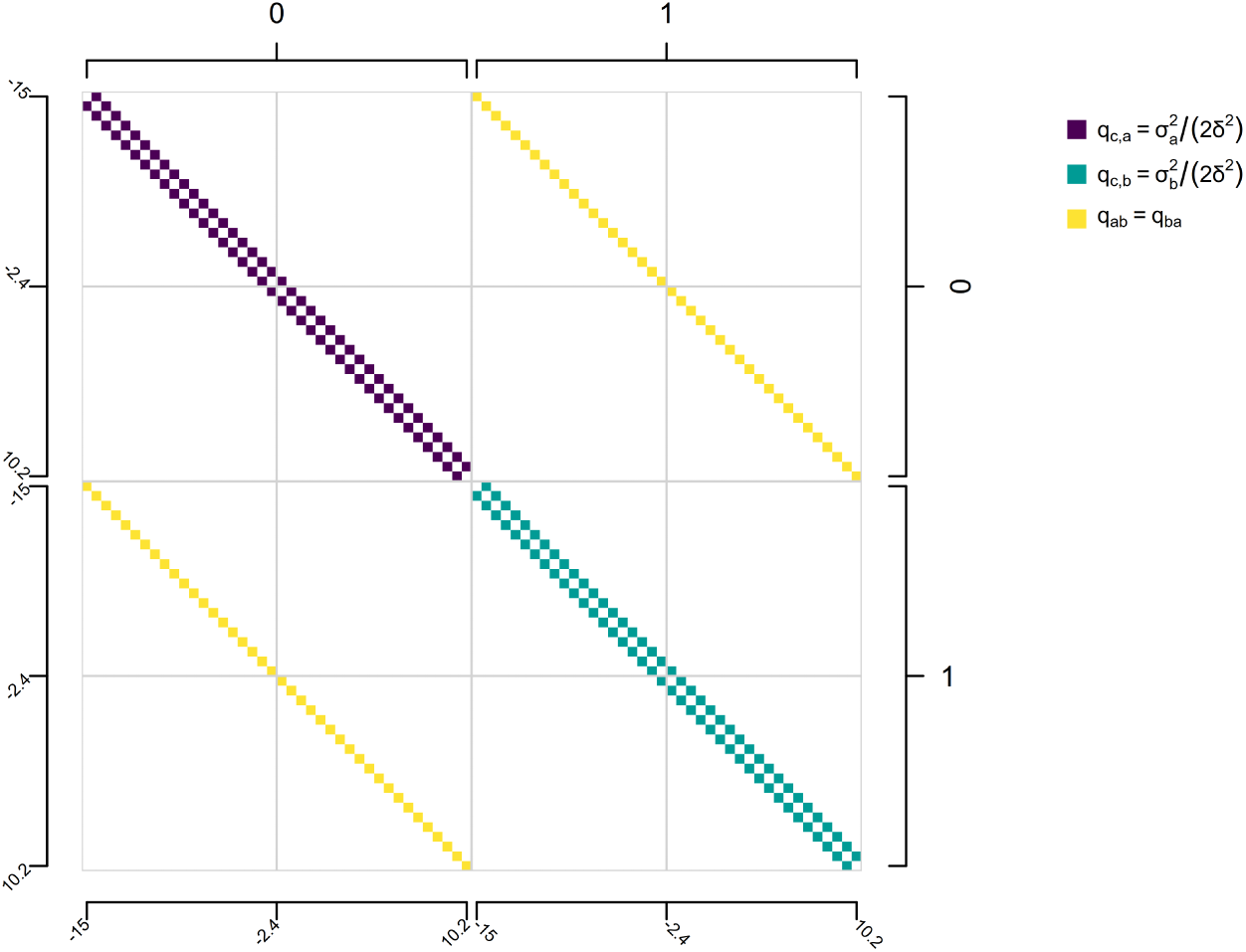
Illustrative figure showing the design of the transition matrix (i.e., infinitesimal generator matrix) of a discrete-state-dependent multi-rate continuous character evolution model for the discretized diffusion approximation. Here, 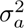 give the discrete character dependent transition rates between adjacent bins for our continuous trait, while *q_ab_* and *q_ba_* give the discrete character transition rates which are here assumed to be equal one to the other, but need not be in general. Merely to make the visualization practical, the number of bins for discretization, *n*, has been set to 40, though in practice a much larger number would be used to obtain accurate parameter estimation and model selection.

Having computed the probability of discretized continuous and discrete character data **x***^′^* using standard pruning (Felsenstein 1981), and just as for our bounded model, above, the probability density of our continuous trait data can be estimated as *f*^^^*_δ_*(**x**) = *p*(**x***^′^*)*/*(*δ^N^*), in which **x***^′^* once again contains the discretized values of our continuous trait *x*. As before, *p*(**x***^′^*) is the probability of the discretized data, *δ* is the continuous character bin width, and *N* is the number of tips in our phylogeny. *f*^^^*_δ_* (**x**) is the discretized diffusion approximation of the probability density function, which is expected to converge to the exact density function (that is, *f*^^^*_δ_* (**x**) *→ f* (**x**)) as *δ →* 0.

To examine the effectiveness of this method in simultaneously estimating the discrete state specific rates of continuous trait evolution, 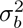 and 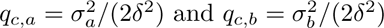, and the rate of transition between discrete states, *q_ab_* and *q_ba_*, we undertook a small simulation experiment.

First, we simulated forty 200-taxon stochastic pure-birth phylogenetic trees with total tree depths of 1.0. On each tree, we next generated a binary discrete character history by first drawing the transition rate between character levels, *q_ab_* = *q_ba_*, at random from a uniform distribution on the interval [1, 10]. This corresponds to quite a high rate of transition between character states, given our total tree depth, which was done on purpose because if the rate of transition is low, then our discrete character changes very seldom and we expect the distinction between our new joint vs. the traditional two-step estimation methods to become much less significant. Then, we simulated continuous trait evolution that depended on our discrete trait histories, with 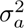 drawn randomly from a uniform distribution on [0.5, 2] and 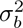 drawn randomly on [5, 20]. This ensures that 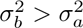, though the magnitude of difference between the two rates varied widely across simulations.

(Readers might note that the *scale* of this simulation study is smaller than we’ve used for previous models. This is because estimation is inherently more computationally challenging for this model than for the others described to this point. The reason for this is largely beyond the scope of the present article, but pertains primarily to the difficulty of large matrix exponentiation. This problem will become even more substantive for the final two models of this article, to be discussed below.)

Having simulated our data in this way, we proceeded to fit our discrete-trait-dependent multi-rate Brownian motion model using the discretized diffusion approximation of Boucher and Démery (2016) and this article. Parameter estimates for 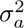 and 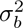 compared to their generating values are given in Figures 12a and 12b. In general, we found that this procedure resulted in unbiased and reasonably accurate estimates of both evolutionary rates in the model, 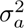 and 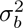 (Figures 12a, 12b).

**Figure 12:**
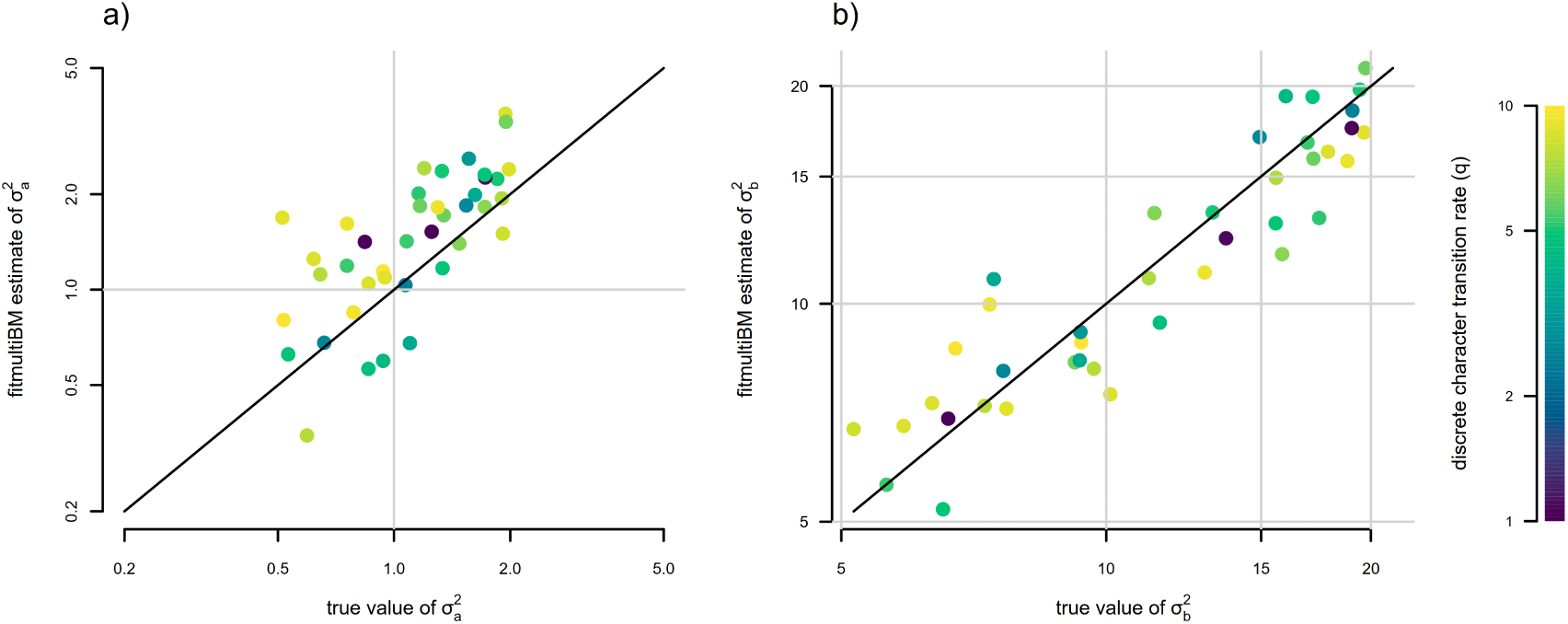
Parameter estimation under a discrete character dependent multi-rate continuous trait evolution model obtained using the discretized diffusion approximation. a) Generating vs. estimated 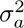. b) Generating vs. estimated 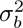. The color gradient of points matches the generating transition rate between discrete character states *q_ab_* = *q_ba_*, with lighter colors corresponding to higher transition rates for the discrete trait. Model parameter estimates were obtained using the *phytools* R package (Revell 2024). See main text for additional details.

We also elected to assess estimation of the rate of transition between discrete character levels in the fitted model, *q_ab_* = *q_ba_*. We found that estimation in the discrete state dependent model was unbiased (Figure 13a). When we compared joint estimation of *q_ab_* = *q_ba_*to estimation using a standard M*k* model (Figure 13b), we also found that the former was slightly more accurate than the latter. In particular, across our 40 simulations, the correlation between estimated and generating values of *q_ab_* = *q_ba_*was *r* = 0.81 using joint estimation via the discretized diffusion approximation, while *r* = 0.74 when *q_ab_* = *q_ba_* estimates were obtained by fitting a standard M*k* model (Figures 13a, 13b).

**Figure 13:**
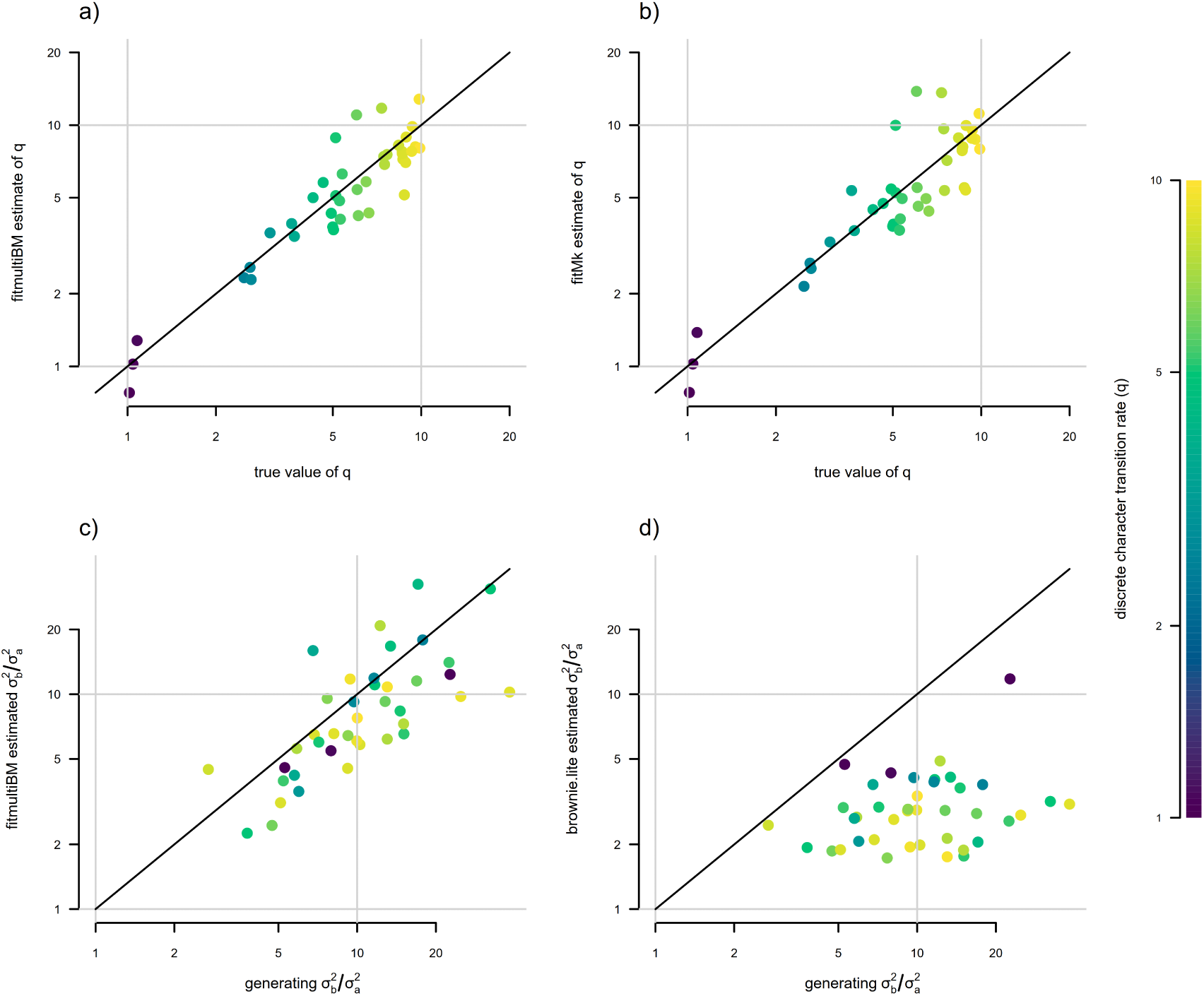
Parameter estimation for joint vs. separate modeling of discrete and continuous character evolution, under a generating scenario of discrete state dependent multi-rate Brownian motion evolution of the continuous trait. a) Generating vs. estimated values for the transition rates between discrete character levels *a* and *b* of the discrete trait, *q_ab_* = *q_ba_*, obtained from the joint model. b) Generating vs. estimated *q_ab_* = *q_ba_* obtained from a standard M*k* model. In these simulations joint modeling was slightly more accurate (*r* = 0.81 compared to *r* = 0.74). c) Generating vs. estimated 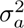 from the joint model. d) Generating vs. estimated 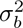 for a two-step procedure, in which 100 stochastic character maps of the discrete character were first obtained, and then a state-dependent continuous trait evolution model was fit to each one and then averaged. Separate modeling of discrete and continuous trait evolution can result in a substantive downward bias in 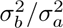, as previously shown in Revell (2013). The color gradient of points corresponds to the generating tra nsition rate between discrete character states, *q_ab_* = *q_ba_*, with lighter colors matching higher generating transition rates, as indicated in the right figure legend. Note that lighter points tend to correspond with greater downward bias in 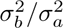 in panel (d), while no such trend is apparent under joint estimation (c). Model parameter estimates w ere obtained using the *phytools* R package (Revell 2024). See main text for more details.

In addition to this analysis, we also elected to compare estimation of 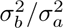 and 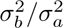 between jointly modeling discrete and continuous trait evolution, and the traditional two-step procedure of first undertaking stochastic

character mapping following Huelsenbeck et al. (2003), fitting a state-dependent rate heterogeneous trait evolution model to each stochastic character map using the method of O’Meara et al. (2006), and then averaging across stochastic maps (Revell and Harmon 2022). Rather than directly comparing parameter estimates themselves, we compared the correlation of the ratio of 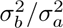 between generating values and their estimates via these two contrasting methods (Revell 2013). The results from this analysis are shown in Figures 13c and 13d. In general, we found that 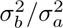 was nearly unbiased in our joint model fit using the discretized diffusion approximation across a range of generating 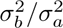 varying from 3 to almost 40 (Figure 13c). By contrast, estimated 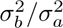 was strongly *downwardly* biased when obtained using the traditional two-step procedure (Figure 13d), particularly for higher transition rates of the discrete character, *q_ab_* = *q_ba_*. This reiterates the general finding of Revell (2013).

In summary, in this subsection we’ve presented a trait evolution model in which a discrete character evolves under a standard continuous time Markov chain (i.e., M*k* model), but the Brownian motion rate of evolution of a separate continuous trait depends on the state of this discrete phenotypic characteristic. This is a common hypothesis in phylogenetic comparative research, and a standard workflow would be to first sample stochastic character map histories of the discrete trait (Huelsenbeck et al. 2003), and then fit a multi-rate Brownian model to each stochastic map (O’Meara et al. 2006; Revell and Harmon 2022). We show how this model can be fit explicitly using the discretized diffusion approximation, as well as how so doing results in less biased estimates of the difference in rates between different discrete character regimes (Revell 2013).

### 2.4 A continuous-character-dependent discrete trait evolution model

In addition to a model in which the state of a discrete character influences the stochastic diffusion of our continuous trait, which we saw in the previous section, the discretized diffusion approximation also permits us to quite easily imagine and fit the inverse model where the state of a continuous character influences the transition rate of a discrete trait. A biological scenario in which we might hypothesize that this model could be a good explanation for our data might include one where the rate of transition to a particular trait value, like viviparity or live birth in reptiles (from egg-laying), is influenced by a continuously-valued environmental factor, such as temperature (e.g., Shine 1983; Lambert and Wiens 2013; Pyron and Burbrink 2014). Lots of other conceivable hypotheses of discrete phenotypic trait evolution could imply a similar relation.

Figures 14c and 14d illustrate the general structure of this process in which we’ve supposed a sigmoidal relationship between the value of our continuous trait and the rates of substitution between the two levels of a continuous character dependent discrete trait, *a* and *b*. This is an arbitrary decision, and our general approach should (in principle) work for any functional form for the relationship between our continuous character and the transition rates of the discrete trait. The sigmoid form is convenient because (depending on its parameterization) it can be designed to be positive everywhere and includes continuous character independence (i.e., a “flat” curve) as a special case. Nonetheless, it would be straightforward to adapt our implementation of this method to other functional forms. As far as we know, and like the semi-threshold model discussed in a previous section, this model is novel to the present article (but see Ives and Garland 2010 or Hadfield and Nakagawa 2010 for statistical approaches designed to investigate a similar underlying biological hypothesis).

**Figure 14:**
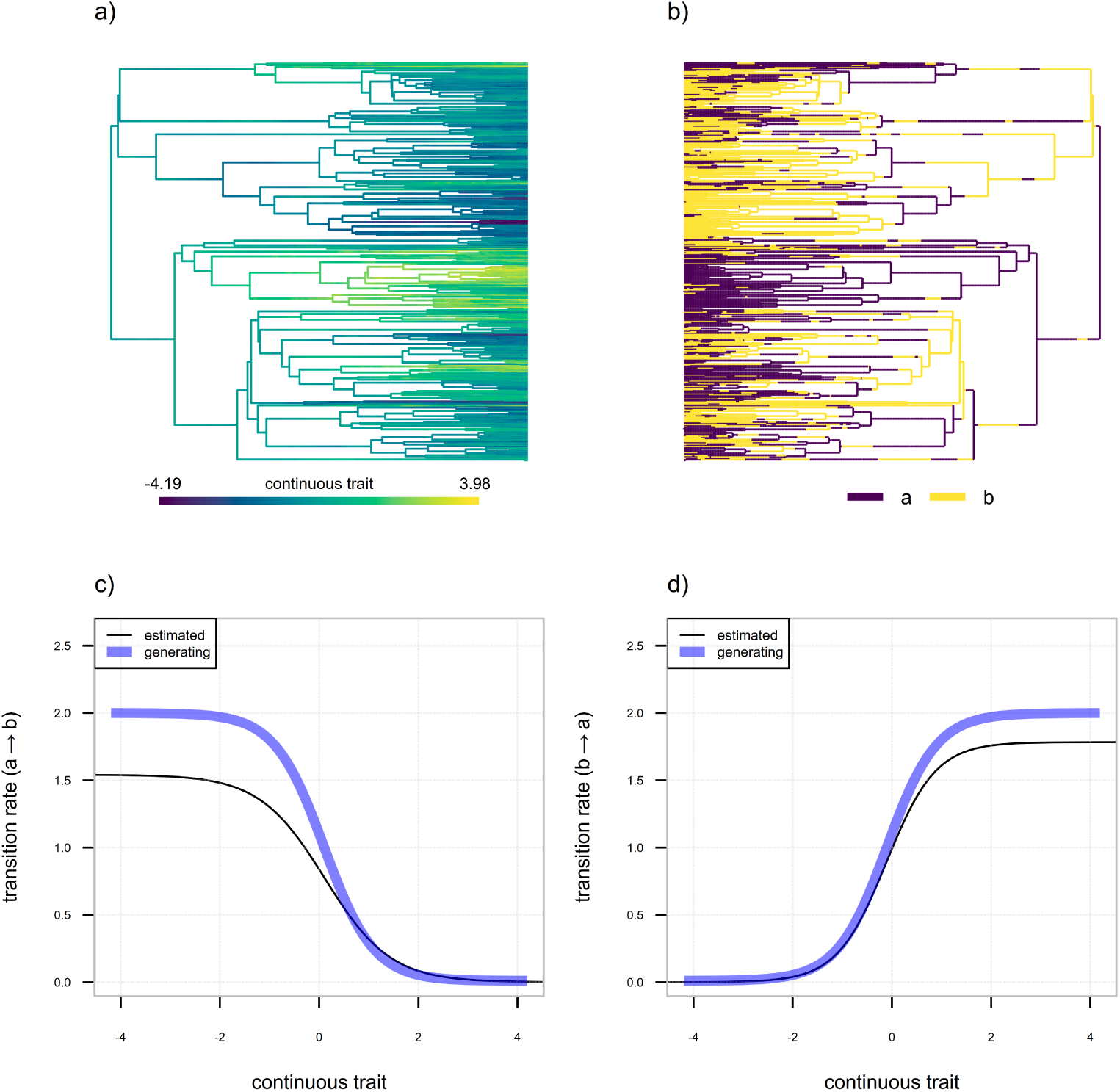
Illustration of the evolutionary process for a binary discrete trait whose rate of evolution depends on the value of a continuous trait. a) Evolutionary history of a continuous trait evolving by Brownian motion on the tree. b) Evolutionary history of a binary discrete character with two levels, *a* and *b*, whose rates of transition between states depend (according to different functional forms in both directions) on the value of the continuous trait of panel (a). c) Generating (blue line) and estimated (black line) transition rate from discrete trait level *a* to *b*, *q_ab_*, as a function of the continuous character of panel (a). d) Generating (blue line) and estimated (black line) transition rate from trait level *b* to *a*, *q_ba_*, as a function of the continuous character value.

In the scenario illustrated by the figure, the “forward” transition rate of our discrete character (*q_ab_*) is high for low values of our continuous trait and low for higher values, while the converse is true with respect to the “backward” transition rate from *b* to *a*, *q_ba_* (Figures 14c, 14d). Data for a continuous character and discrete trait simulated on a stochastic phylogeny with 400 tips under these conditions is illustrated by Figures 14a and 14b, respectively. In the figure, we see that the evolutionary process has predictably resulted in an association between relatively high values of our continuous character (Figure 14a) and condition *b* of the discrete trait (Figure 14b), as well as between relatively low values of the continuous character (Figure 14a) and condition *a* of the discrete trait (Figure 14b). This is one scenario of continuous character trait-dependent discrete character evolution; however, our model would also be appropriate for conditions in which (for instance) one extreme of the continuous trait distribution was associated with both higher backward and forward rates of the discrete character, and the other with low discrete trait rates. Though this evolutionary process unambiguously corresponds to continuous-character-dependent rate-varying discrete trait evolution, it would not tend to result in the same predictable character association shown by Figures 14a and 14b (also see Maddison and FitzJohn 2014). It would nonetheless be appropriate to study using our new model.

To fit this model to data using the discretized diffusion approximation, we use an infinitesimal generator (transition matrix) of our discretized space that is structurally similar to that of Figure 11, but wherein the transition rates, *q_ab_*and *q_ba_*, are allowed to vary as a continuous function of our quantitative trait, rather than assumed to take a constant value. As previously noted, in our implementation we suppose a sigmoidal functional form for this relationship, but there’s no inherent necessity that it take this specific shape. Additionally, and unlike the generator matrix of Figure 11, here we assume a constant *σ*^2^ for the continuous trait (though this assumption, too, could easily be relaxed). The design of this generator matrix is illustrated in Figure 15. As in Figure 11, this matrix is of dimensionality *k · n × k · n*, for *k* levels of our discrete trait (where *k* = 2 in this case) and *n* bins of discretization for the continuous character in our model.

**Figure 15:**
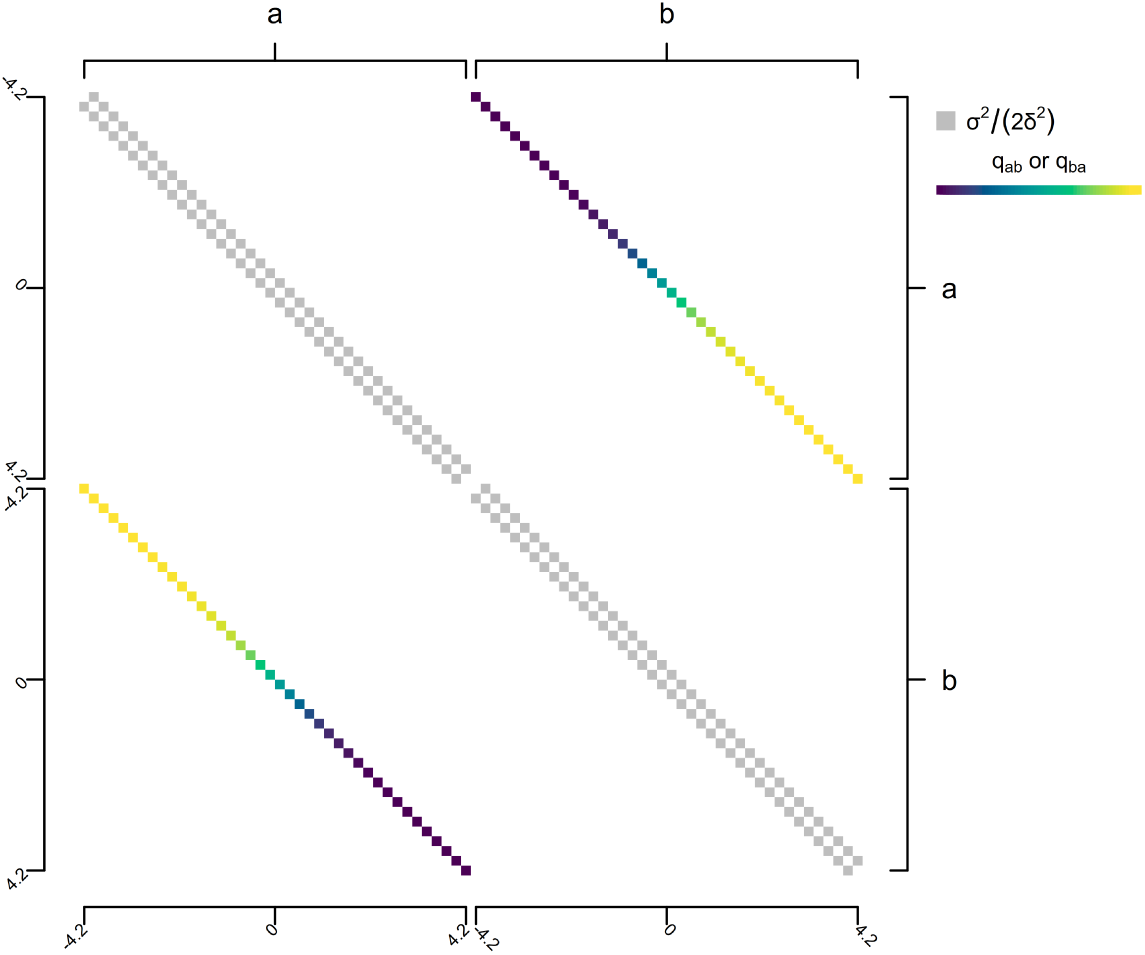
Illustrative figure showing the design of the infinitesimal generator (transition matrix) of a continuous-character-dependent multi-rate discrete character evolution model for the discretized diffusion approximation. Here a specific sigmoidal form is being used to match the generating conditions of Figure 14, though in practice this form is precisely what would be estimated from the data. For visualization purposes only, the number of bins for discretization has been set to 40. In practice a much larger number would be used to obtain accurate parameter estimation and model selection. See main text for more details.

Unfortunately, for computational reasons beyond the scope of the present study (but that once again pertain to large matrix exponentiation and that we hope to overcome via future research), it was not feasible for us to undertake even a modestly-sized simulation experiment to assess estimation under this model. We have nonetheless provided the maximum likelihood fitted model for the data and tree of Figures 14a and 14b in the solid black lines of Figures 14c and 14d. Note that all parameters of the backward and forward transition rate functional forms are being estimated from the data and phylogeny: the lower and upper asymptotes, the mid-or inflection point, and the curve steepness (Figure 14). At least in this single simulated example, the similarity of our estimate to the generating conditions is notable (Figures 14c, 14d). Readers interested in employing this method for their own research are encouraged to download our simulation and analysis code (which is publicly available, see below) and undertake their own study of parameter estimation reliability for this model.

In summary, we have presented a new model for the joint evolution of a single discrete and separate continuous character in which the forward and backward transition rates of the discrete trait depend on the value of the continuous trait according to sigmoidal functional forms that are free to vary in the two directions of discrete character transition (Figure 14). We show how this model can be fit to data using the discretized diffusion approximation. For computational reasons, we do not provide an extensive simulation study of this method; however, we recommend that it be investigated by future research.

### 2.5 A multi-trend directional model

As a final illustration of the power of our discretized diffusion approximation, we’ve chosen to develop another new model: a discrete-character-dependent multi-trend (i.e., multi-*µ*), multi-rate (i.e., multi-*σ*^2^), trended or directional Brownian motion model. This is a discrete-character-dependent stochastic diffusion model, but where diffusion has a linear trend (*µ*) and in which both the diffusion rate (*σ*^2^) and the trend (*µ*) vary as a function of our discrete trait. This evolutionary scenario is illustrated via the simulation of Figure 16. Note that we intentionally simulated a non-ultrametric tree because the trend model parameter *µ* and *x*_0_ are mutually non-identifiable (that is, infinitely many *µ* and *x*_0_ pairs result in the same likelihood) for ultrametric phylogenies (e.g., Ho and Ané 2014b; Gill et al. 2017).

**Figure 16:**
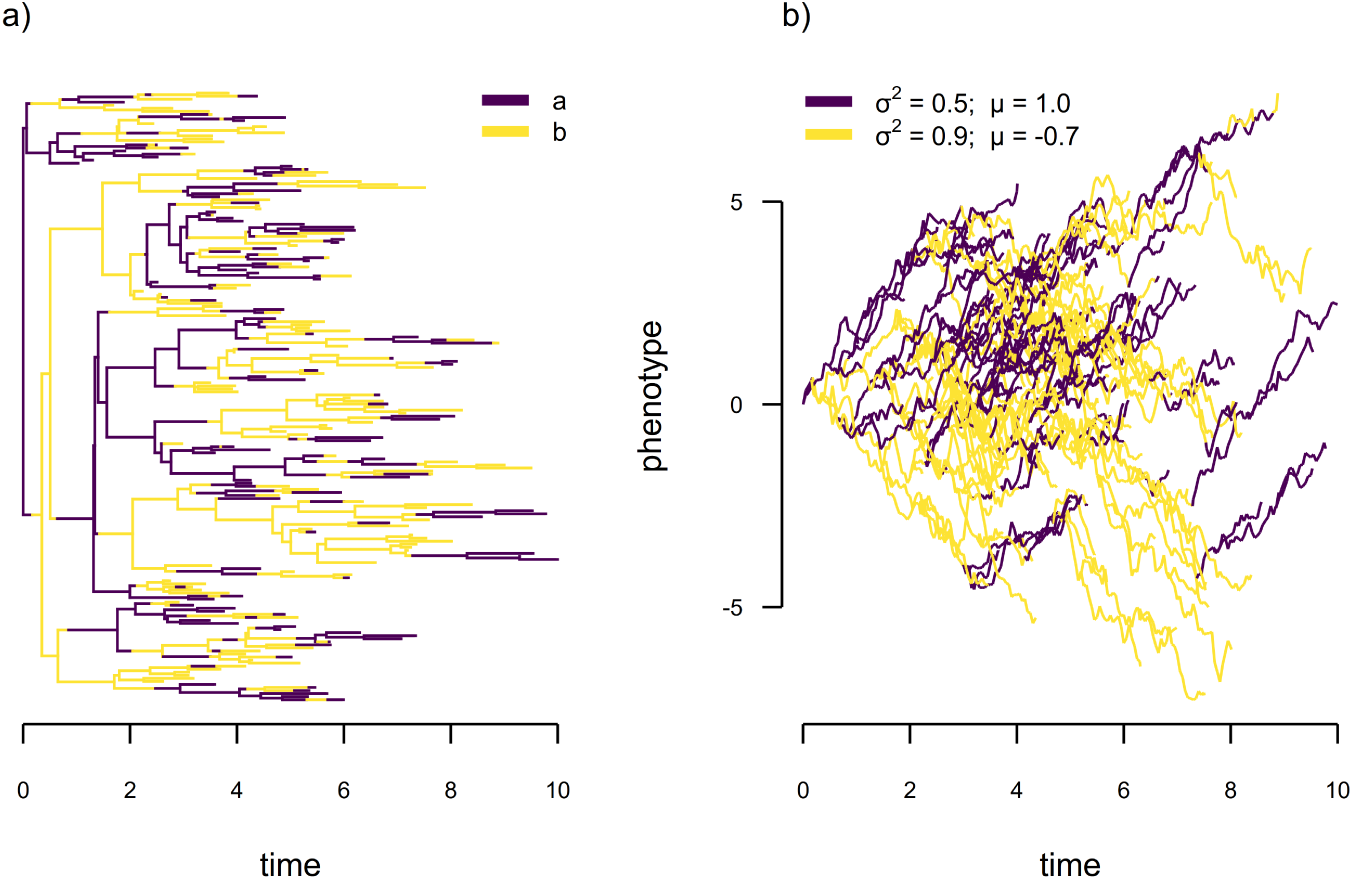
Illustration of the scenario of discrete-character-dependent multi-trend, multi-rate trended Brownian motion evolution. a) Simulated discrete character evolution of a binary character with two levels, *a* and *b*, where *q_ab_*= *q_ba_* = 0.5. b) Continuous trait evolution under our model for 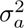 = 0.5, 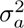 = 0.9, *µ_a_*= 1.0, and *µ_b_* = *−*0.7. Under this scenario, the linear trend of our continuous trait is positive (i.e., increasing) when the discrete character is in condition *a* and negative (decreasing) when in condition *b*.

As far as we know, this specific model in phylogenetic comparative methods is novel to the present article; however, Gill et al. (2017) provide a relaxed random walk model in which they also propose to jointly model continuous trait and nucleotide sequence evolution. (This is related to what we have done here, but assumes that the discrete characters and continuous trait evolution are conditionally independent, Gill et al. 2017.) Among many possible applications, a biological scenario for this model might be one in which trended geographic diffusion of a host-switching wildlife virus is expected (e.g., Gill et al. 2017), but where the trend direction switches as a function of the viral host, which in turn evolves via a continuous-time Markov chain.

To fit this model to data using the discretized diffusion approximation we take the same approach as for the preceding two models; however, here our diffusion process is asymmetric in the upward and downward directions. That is to say, *q*_+_ = *σ*^2^*/*(2*δ*^2^) + *µ/*(2*δ*) and *q_−_* = *σ*^2^*/*(2*δ*^2^) *− µ/*(2*δ*) (where *q*_+_ and *q_−_* refer to the upward and downward transition rates between adjacent bins of our discretized continuous trait space for a given set of *σ*^2^ and *µ* values) for diffusion rate *σ*^2^ and linear trend parameter *µ*, where *δ* is as previously defined (Risken 1996; Kushner and Dupuis 2001). (Since both *q*_+_ and *q_−_* must be *≥* 0, fitting this model using the discretized diffusion approximation also requires that we select *δ* such that *|µ| ≤ σ*^2^*/δ*.) The macrostructure of the transition matrix for this model is identical to that illustrated in Figure 11 except that the super-and sub-diagonals of each major axis submatrix are allowed to assume different values as a function of the trend, *µ*, of each discrete character level.

To illustrate this model, we undertook a limited simulation study. We simulated twenty 200-taxon non-ultrametric trees of total depth 10.0, and a binary discrete trait with *q_ab_* = *q_ba_*= 0.5, as in Figure 16. We then sampled *µ_a_* and *µ_b_* randomly on [0.2, 1] and [*−*1*, −*0.2], respectively. This way the trend, *µ*, was always positive for discrete character level *a* and negative for *b*. We randomly sampled 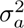 and 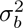 independently on [0 3 1]. For each tree and dataset we fit the multi-μ, multi-*σ*^2^ model using the discretized diffusion approximation.

In general, we found reliable estimation of the simulated discrete character dependent stochastic diffusion rates (*σ*^2^ and *σ*^2^) and linear trend parameters (*µ_a_* and *µ_b_*) in this model (Figure 17), though many more simulations would be required to truly understand its properties.

**Figure 17:**
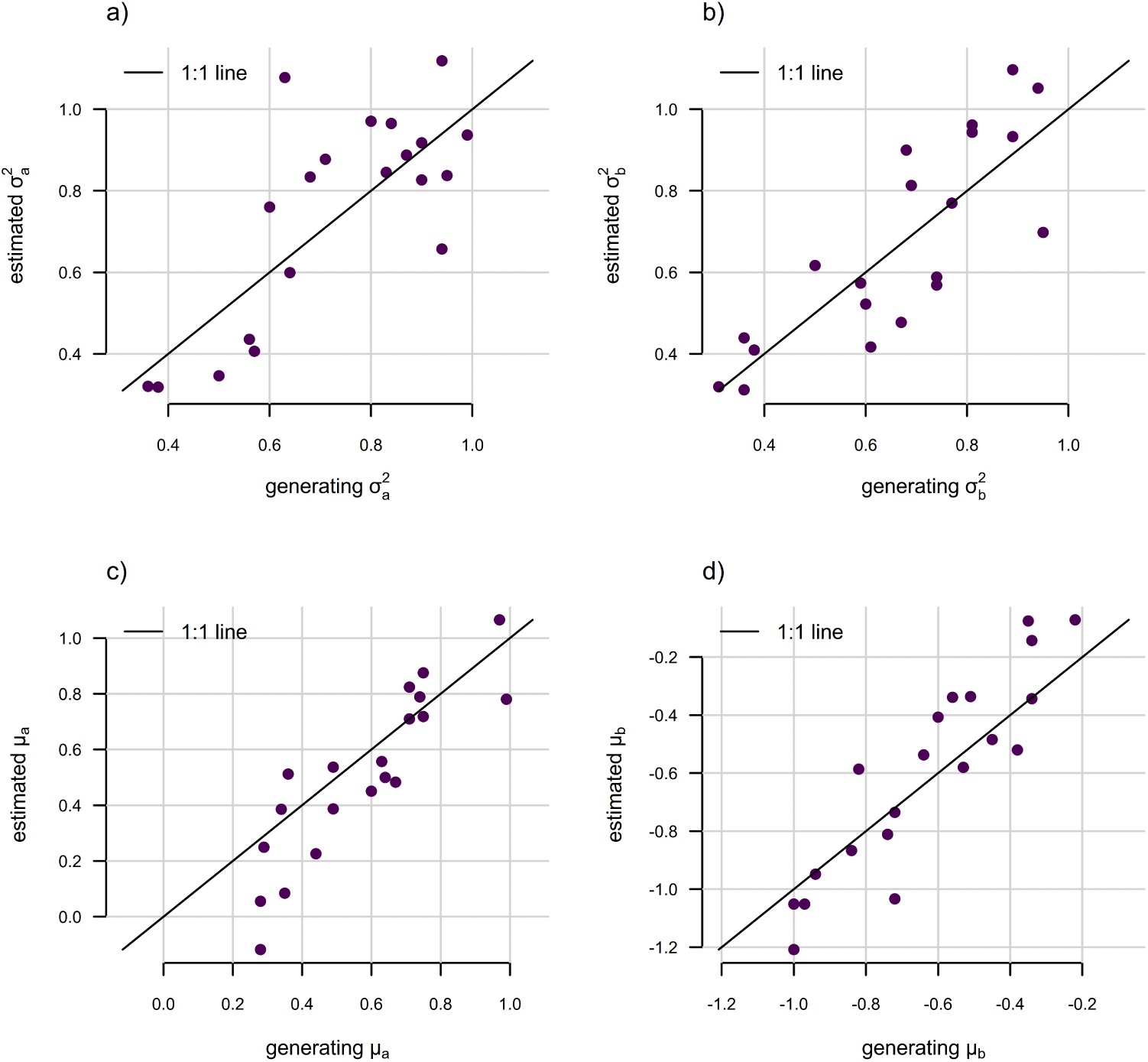
Parameter estimation under discrete-character-dependent multi-trend trended Brownian motion evolution. The discrete character was simulated with two levels, *a* and *b*, for a constant transition rate between levels, *q_ab_* = *q_ba_* = 0.5. a) Generating vs. estimated 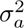 under the model. b) Generating vs. estimated 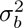. c) Generating vs. estimated *µ*, the linear trend parameter of the model when the discrete trait is in condition *a*. d) Generating vs. estimated *µ*. Model parameter estimates were obtained using the *phytools* R package (Revell 2024). See main text for additional details.

We also checked parameter estimation for the rate of transition of the discrete trait in the model, *q_ab_* = *q_ba_*. We found no difference between *q̂* in our fitted trend models (*q̅* = 0.47 compared to a constant true value of *q* = 0.5) and a standard M*k* model (*q̅* = 0.48), although the former had a slightly lower variance across simulations than the latter (*σ* = 0.076 for joint estimation compared to 0.10 for the standard M*k* model). Given the small scope of simulations this result should be considered very preliminary; however, it comports with the finding of greater accuracy in the estimation of *q_ab_* = *q_ba_* we reported for the discrete-character-dependent rate-varying Brownian model earlier in this article (Figure 13).

In summary, we present a new discrete character dependent multi-trend trended Brownian motion model. We describe how this model can be fit to data using the discretized diffusion approximation. We anticipate that the model will only be fully identifiable for non-ultrametric phylogenetic trees, and show via a very limited simulation study that it seems to have reasonably good statistical properties.

## 3 Software implementation and data availability

All of the methods of this article are implemented in the R statistical computing software (R Core Team 2025) package *phytools* (Revell 2012, 2024). All analyses of this article were conducted using *phytools*, *geiger* (Harmon et al. 2008; Pennell et al. 2014), and *ape* (Paradis and Schliep 2019). *phytools* and *geiger* also in turn depend on the core R phylogenetics packages *ape* (Paradis et al. 2004; Paradis and Schliep 2019) and *phangorn* (Schliep 2011). *phytools*, *geiger*, *ape*, and *phangorn* are all publicly available through the Comprehensive R Archive Network, CRAN (https://cran.r-project.org/). All data and code of this article can be reviewed via a public GitHub repository (https://github.com/liamrevell/Revell_etal.unlocking_pcms).

## 4 Concluding thoughts

A decade ago Boucher and Démery (2016) published an article that one of us (LR) has been consistently asserting since 2024 is “the most important phylogenetic comparative methods paper you probably haven’t read.” This is at least partly because LR either had not read the article when it was published or had read it without properly appreciating it, and did not do so for several years afterward. Though certainly many others read the article, it’s easy to argue that it was underappreciated by the field since it had been cited a mere 55 times since its publication at the time of initial submission of this article. (For reference, this tally almost exactly matches the number of times an LR lab undergraduate-authored paper on lizard tail autotomy in the *Journal of Herpetology* from the same year has been cited as of the time of writing, Tyler et al. 2016.)

The Boucher and Démery (2016) article introduced a new model, bounded Brownian motion, and at the same time circumvented the intractability of the likelihood function for reasonably-sized trees by devising a clever procedure for approximating the likelihood using what we’ve referred to herein as the discretized diffusion approximation. In essence, this approach replaces the underlying diffusion process with a continuous-time Markov chain defined on a discretized trait space with transition rates between adjacent bins chosen so that the chain approximates the generator of the diffusion process. This sort of construction is closely-related to classical finite-difference approximations of diffusion widely employed throughout the stochastic process literature (e.g., Kushner and Dupuis 2001; Gardiner 2009; Pavliotis 2014). What was truly novel about Boucher and Démery (2016), then, was not the general mathematical idea, but its application to phylogenetic comparative models. As discussed in the Introduction, a subsequent contribution by some of the same authors, Boucher et al. (2018), extended this latter innovation to a wider variety of continuous trait evolution models; however, in our opinion, this article is also underrecognized.

In 2024, two of the other authors of the present article (MA and LH) invited the lead author (LR) to participate in a study they were working on (with author KM and others) on the topic of “circular” Brownian motion (Martinet et al. In review), also using the technique of Boucher and Démery (2016). At that time, it became apparent to LR that there might be a substantially larger number of other applications of this estimation procedure than those that had already been identified. This was the origin of the present article.

Herein we’ve presented (as far as we can ascertain from the literature) at least three or four totally novel models for the phylogenetic comparative analysis of discrete and/or continuous trait evolution. Nonetheless, we also confidently assert that this barely scratches the surface of the broad range of potential uses of this powerful approach.

Some additional applications that we have not covered include a wide variety of so-called “hidden-rate” models (Beaulieu et al. 2013; Boyko and Beaulieu 2021). In a hidden-rate model, we might hypothesize discrete-character-dependent continuous trait evolution, but where the discrete character is unobserved, thus allowing the continuous trait data to entirely dictate the phylogenetic position of shifts between rates or evolutionary regimes. Indeed, LR has already implemented precisely this model (in a few flavors) within his R package *phytools* (Revell 2024). Similarly, it’s straightforward to extend the same logic to our continuous-character-dependent rate-varying discrete trait model, though, at the time of writing, this had not yet been implemented in software. In addition to models in which merely the rate or direction of Brownian evolution varies as a function of a discrete trait, the discretized diffusion approximation should also allow us to vary the process of evolution, for example, under an Ornstein-Uhlenbeck diffusion with a “rubber band” process (Hansen 1997; Butler and King 2004). Between initial submission of this article and its present revision, this model, too, was developed and is already available in the *phytools* R package (Revell 2024). As opposed to bounded evolution of Boucher and Démery (2016) or the semi-threshold model of the present article, we also might imagine evolutionary limits in which the bounds were absorbing or “sticky.” This model is straightforward to implement using the discretized diffusion approximation and is already available to both simulate and fit to data in *phytools* (Revell 2024). We’ve focused exclusively on “gradual” continuous character models, but the discretized diffusion approximation would easily extend to process models with discontinuous jumps. For example, it would be very easy to imagine and implement in software a model of continuous character evolution but where a trait is “turned off” or lost in a single evolutionary event, though we have yet to explore this sort of model.

We believe that both what we’ve covered in this article and mentioned above could represent but the tip of a much larger iceberg of evolutionary scenarios for which likelihood functions are intractable but where the discretized diffusion approximation may be suitable. As such, we fully expect to continue discovering new applications of this approach into the future.

The famous Maslow (1966) law of the instrument states “if all you have is a hammer, everything looks like a nail” and is meant as a cautionary tale. The implication is that if you have but a single tool at your disposal, a cognitive distortion can arise in which many different problems will begin to look as if they can be remedied using this instrument. What we seem to have discovered in exploring the broad usefulness of the Boucher and Démery (2016) finite-difference or discretized diffusion approximation for the phylogenetic analysis of discrete and continuous trait data is a sort of inverse law of the instrument. If you look closely enough at many hypotheses and problems in phylogenetic comparative research that pertain to trait evolution, many do, in fact, turn out to be nails.

## Acknowledgments

This project was funded in part by a Tufts IRACDA Partner Faculty Research Award Program grant (NIH-NIGMS award K12GM133314, sub-awarded to LR) and by a Proposal Development Grant from the University of Massachusetts Boston (to LR and NH). We used ChatGPT (OpenAI) for grammatical and typographical proofreading, to check and improve technical clarity, and to assist in literature review; however, no portion of this article was written by AI and all content was reviewed and verified by the authors.

